# Genome-resolved insight into the reservoir of antibiotic resistance genes in an aquatic microbial community

**DOI:** 10.1101/2022.08.30.505784

**Authors:** Zahra Goodarzi, Sedigheh Asad, Maliheh Mehrshad

## Abstract

Aquatic microbial communities are an important reservoir of Antibiotic Resistance Genes. However, distribution and diversity of different ARG categories in environmental microbes with different ecological strategies is not yet well studied. Despite the potential exposure of the southern part of the Caspian Sea to the release of antibiotics, little is known about its natural resistome profile. We used a combination of Hidden Markov model (HMM), homology alignment and a deep learning approach for comprehensive screening of the diversity and distribution of ARGs in the Caspian Sea metagenomes at a genome resolution. Detected ARGs were classified into five antibiotic resistance categories including Prevention of access to target (44%), Modification/protection of targets (30%), Direct modification of antibiotics (22%), Stress resistance (3%), and Metal resistance (1%). The 102 detected ARG containing metagenome-assembled genomes of the Caspian Sea were dominated by representatives of Acidimicrobiia, Gammaproteobacteria and Actinobacteria classes. Comparative analysis revealed that the highly abundant, oligotrophic, and genome streamlined representatives of taxa Acidimicrobiia and Actinobacteria modify the antibiotic’s target via mutation to develop antibiotic resistance rather than carrying extra resistance genes. Our results help with understanding how the encoded resistance categories of each genome are aligned with their ecological strategies.

## Introduction

Antibiotic resistance is a major threat to modern society. Projections indicate that the antimicrobial resistance (AMR) attributable mortality could reach up to 10 million by 2050 (1). To tackle AMR crisis as a complex, transboundary, and multifactorial health challenge, understanding the connections between the human, animal and environmental microbiome (the One Health concept) is critical. The dynamics and evolution of AMR depend on the communication networks in local (One Health) and global (Global Health) levels. So, prevention, surveillance and control of AMR require integrated political and socio-economic actions at the global stage and these actions require a comprehensive ecological surveillance networks (2).

Despite its adverse effect on human health, AMR is a natural phenomenon. While it is clear that excessive use of antibiotics significantly contributes to the emergence of resistant strains, antibiotic resistance also exists in natural bacteria of pristine ecosystems (3). Antibiotics and antibiotic resistance genes (ARGs) have been co-evolving in the ecosystems for millions of years (4) (Mostly fueled by microbe’s continuous competition for resources). In addition to their well-known role, antibiotics and ARGs play other physiological roles in nature. For example, at sub-inhibitory concentrations, antibiotics act as signaling molecules involved in quorum sensing and biofilm formation (5,6). Some ARGs were originally involved in cellular functions such as virulence, cell homeostasis and intercellular signal trafficking (7), but were selected for the resistance phenotype and got transferred from the environmental reservoirs into commensal and pathogenic bacteria. Following the widespread presence of antibiotics, this transfer occurred very rapid on an evolutionary scale through horizontal gene transfer (HGT) and mobile genetic elements (MGEs)(8,9). Environmental microbiome have been shown to serve as potential reservoirs of antibiotic resistance genes primed for exchange with pathogenic bacteria (10). Nevertheless, the evolution and prevalence of ARGs in environmental microorganisms is poorly understood (4).

Antibiotics are currently used widely, not just for the treatment of human infections, but also in agriculture, livestock, and aquaculture industries. Discharge of antimicrobials and resistant micro-organisms in waste from healthcare facilities, pharmaceutical manufacturing facilities and other industries into the environment affects the natural ecosystems (11). This has been shown to accelerate development and transfer of AMR among bacterial populations in clinical and natural environments through selection pressures (12). This concern is also growing by global warming as it might accelerate the spread of antibiotic resistance (13).

Meta-omics studies from different natural ecosystems specially aquatic environments such as ocean (14,15), rivers (16), lakes (17) and sea water (18) have recently detected ARGs and profiled the antibiotic resistome of these ecosystems. These studies reiterate that even natural environments that have not been exposed to high antibiotic concentrations could potentially be a reservoir of ARGs. While recovered ARGs in oceanic ecosystems mainly belong to representatives of Gamma- and Alpha-proteobacteria (19) (14), comparative analysis of ARGs present in different taxa in relation to their ecological strategies is missing. Investigating the environmental reservoirs of ARGs, their presence on horizontally transferable mobile genetic elements (MGEs), taxonomic affiliation of the Antibiotic resistant bacteria (ARBs), and their ecological strategies is critical to assess their contribution to emergence and spread of ARGs as well as future actions to fight resistant infections. To this end, we performed genome resolved metagenomic analysis for ARG annotation in the deeply sequenced depth profile metagenomes of the Caspian Sea. The southern part of the Caspian Sea is increasingly exposed to human pollutions. There is a high percentage of organic matter entering the basin via agricultural and aquaculture effluents (20). The status of antibiotic pollution of the Caspian Sea has not been assessed, however as the WHO report on surveillance of antibiotic consumption puts Iran among countries with high-level use of antibiotics (21), monitoring the ARG reservoir of the Caspian sea seems critical. We applied Hidden Markov model (HMM), homology alignment and a deep learning approach supplemented with manual curation of potential ARGs and classified them into five antibiotic resistance categories. These approved ARGs were studied in relation to the ecological strategies of ARBs containing them. The results of metagenomics ARG surveys, are constrained by the comprehensiveness and quality of the used antimicrobial resistance gene databases (22). Here we used different databases of protein and nucleotide sequences as well as different approaches to provide a comprehensive genome-resolved view of the Caspian Sea resistome. Our results show that most of the ARG containing metagenome-assembled genomes (MAGs) are among taxa that are still evading the bound of culture. More interestingly we see that the streamlined genomes mainly contain ARGs with mutations in the antibiotic target rather than carrying extra genes for antibiotic resistance.

## Results and discussion

### Caspian Sea MAGs characteristics

In this study, we explored the diversity and distribution of ARGs in three metagenomes collected along the depth profile of the brackish Caspian Sea. Binning resulted in 477 metagenome-assembled genomes (MAGs) with completeness ≥ 40% and contaminations ≤ 5%. Only 14 MAGs belonged to domain Archaea and the rest of 463 bacterial MAGs were dominated by Proteobacteria, Bacteroidota, and Actinobacteriota (Overall taxonomic distribution in **Supplementary Figure S1**.).

### Antibiotic resistance gene profile of the Caspian Sea Bacteria

Using six different tools we initially detected in total 259 potential ARGs in 110 MAGs. All predicted genes were further checked for conserved domains manually to confirm the functional predictions. For detected genes that confer resistance to antibiotics due to mutations we manually checked the alignments and report them as potential resistance genes only when they contain the exact mutation as those reported to cause resistance. A total of 82 genes conferring antibiotic resistance due to mutation were initially detected. Multiple sequence alignment together with reference genes confirmed mutation in 56 genes, two parC gene, three murA genes, 31 rpsL genes and 20 rpoB genes (multiple sequence alignments and mutations conferring antibiotic resistance are shown in the **Supplementary Figure S2**). There are ongoing debates regarding the relevance of detected mutations in similar genes to exhibiting resistant phenotype in the organism (23–25). While our alignment results suggest antibiotic resistance, further experimental tests are needed to confirm the resistant phenotype.

After this curation step in total 33, 95, and 105 predicted open reading frames (ORF) were identified, and their annotation was confirmed as putative ARGs in respectively 15, 40, and 150 m depth MAGs (Fig. 1d). These ARGs were distributed in 102 bacterial genomes (Fig. 1a). Confirmed ARGs identified via each screening tool are detailed in the **Supplementary Table S1**. Antibiotic resistant bacteria (ARB) were more diverse in 150 m (13 different classes) and 40 m (12 different classes) depths as compared to the 15 m (6 different classes) depth. In general, a higher phylogenetic diversity was detected in the deeper strata of the Caspian Sea (26).

**Figure 1.**
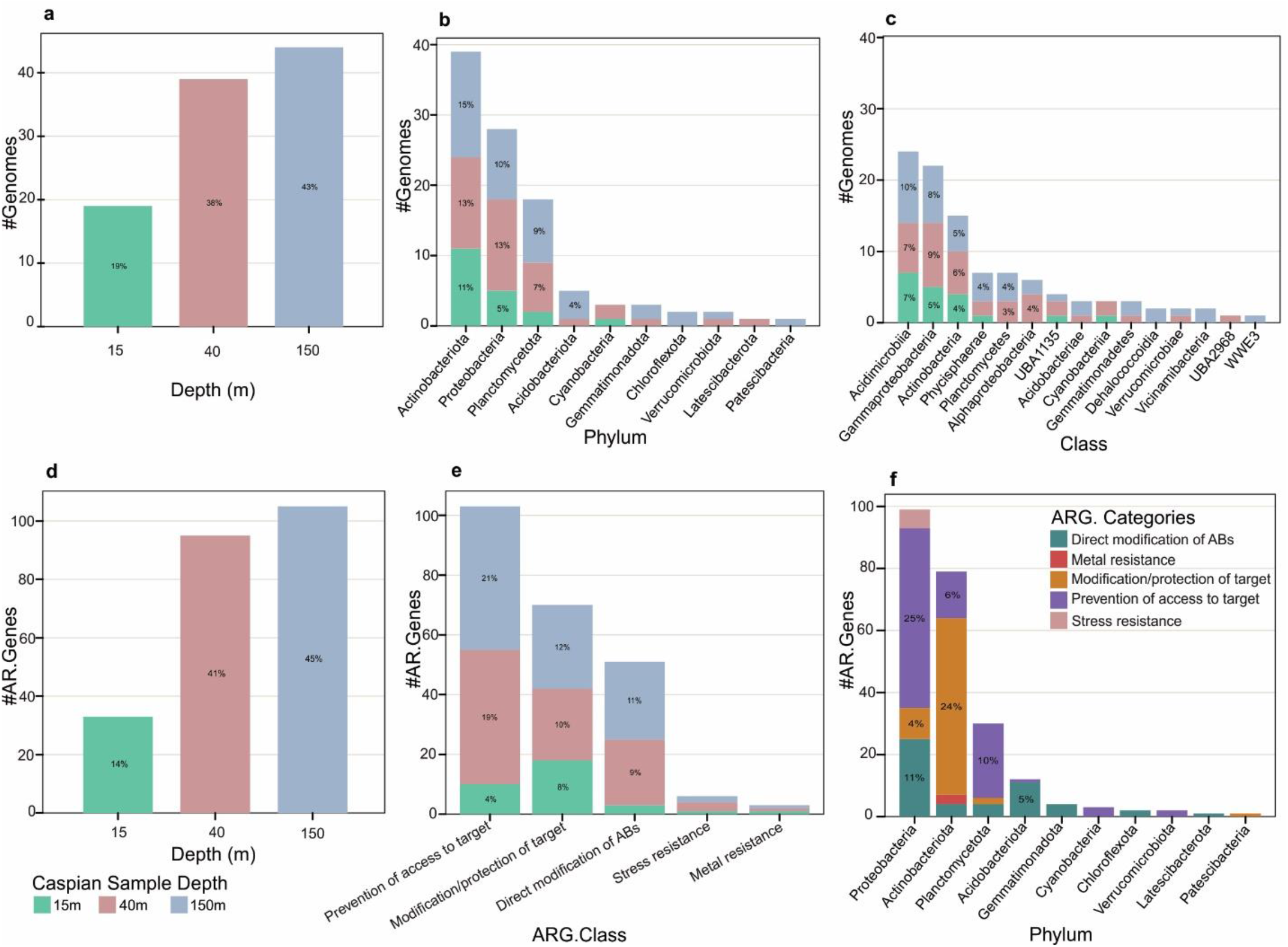
Distribution of ARGs and ARG containing MAGs in the Caspian Sea metagenomes. Distribution of ARG containing MAGs based on depth (**a**), and phylogenetic diversity at the phylum (**b**), and class (**c**) level. Distribution of ARGs in different depth (**d**), abundance of ARGs per ARG categories in different depth (**e**), and abundance of ARG categories based on phylogenetic diversity at the phylum level (**f**).

The Caspian ARGs were classified into 5 different antibiotic resistance categories according to annotated functions: (I) Prevention of access to target, (II) Modification/protection of targets, (III) Direct modification of antibiotics, (IV) Stress resistance, and (V) Metal resistance (stats of these categories and their subcategories are shown in the **Table 1**). The most frequently detected category was prevention of access to target (44%) as a result of antibiotic efflux pumps followed by, Modification and protection of targets (30%) (Fig. 2) and direct modification of antibiotics (22%). The rest of identified ARGs were classified in two categories of Stress (6 genes or 3%) and Metal (3 genes or 1%) resistance. Categories (I), (II) and (III) ARGs were less prevalent in 15 m depth metagenome of the Caspian Sea (Fig. 1e).

**Table 1.**
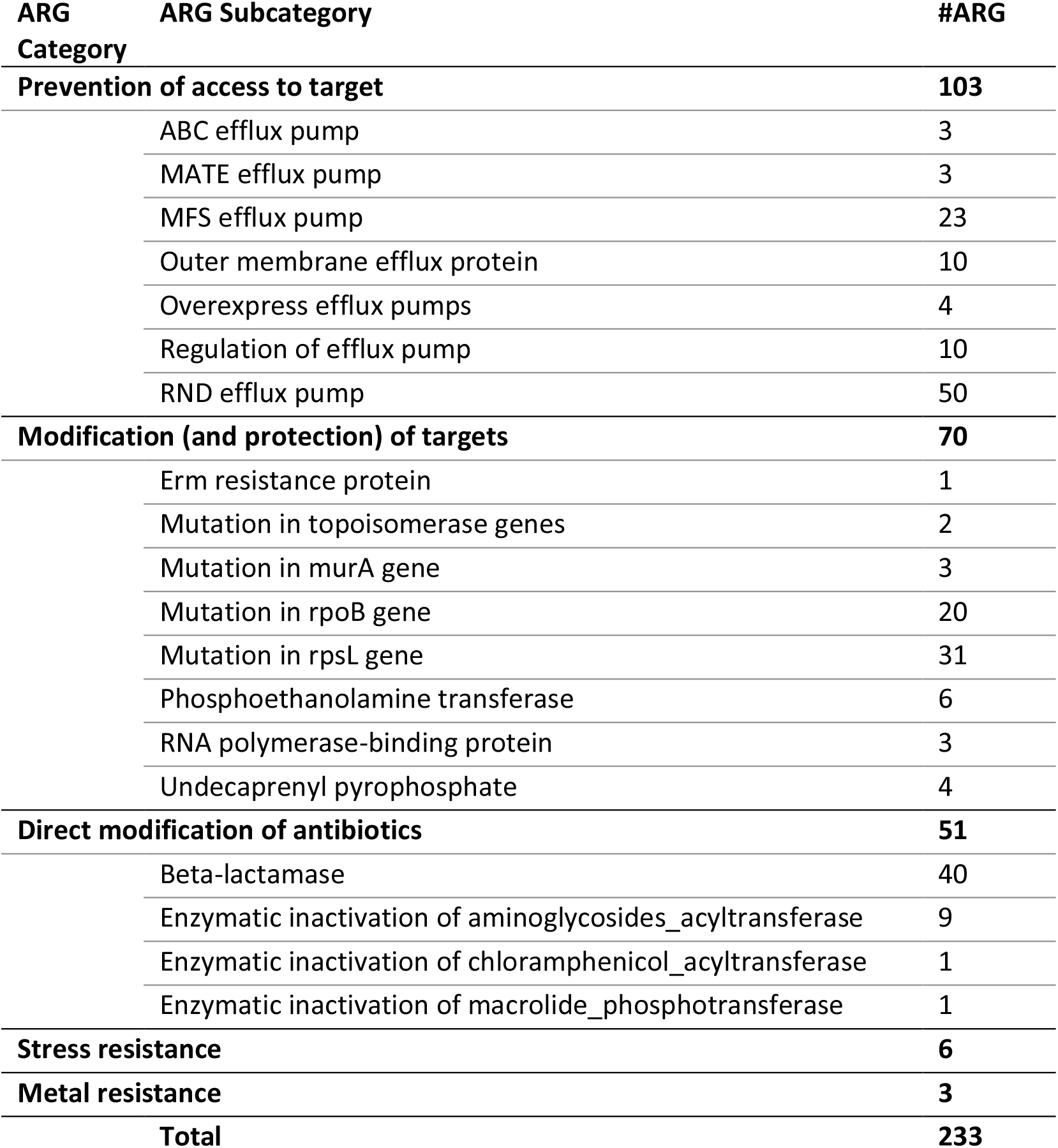
Distribution of detected Caspian Sea ARGs in different categories and subcategories.

| ARG Category | ARG Subcategory | #ARG |
| --- | --- | --- |
| **Prevention of access to target** |  | **103** |
|  | ABC efflux pump | 3 |
|  | MATE efflux pump | 3 |
|  | MFS efflux pump | 23 |
|  | Outer membrane efflux protein | 10 |
|  | Overexpress efflux pumps | 4 |
|  | Regulation of efflux pump | 10 |
|  | RND efflux pump | 50 |
| **Modification (and protection) of targets** |  | **70** |
|  | Erm resistance protein | 1 |
|  | Mutation in topoisomerase genes | 2 |
|  | Mutation in murA gene | 3 |
|  | Mutation in rpoB gene | 20 |
|  | Mutation in rpsL gene | 31 |
|  | Phosphoethanolamine transferase | 6 |
|  | RNA polymerase-binding protein | 3 |
|  | Undecaprenyl pyrophosphate | 4 |
| **Direct modification of antibiotics** |  | **51** |
|  | Beta-lactamase | 40 |
|  | Enzymatic inactivation of aminoglycosides_acyltransferase | 9 |
|  | Enzymatic inactivation of chloramphenicol_acyltransferase | 1 |
|  | Enzymatic inactivation of macrolide_phosphotransferase | 1 |
| **Stress resistance** |  | **6** |
| **Metal resistance** |  | **3** |
| **Total** |  | **233** |

**Figure 2.**
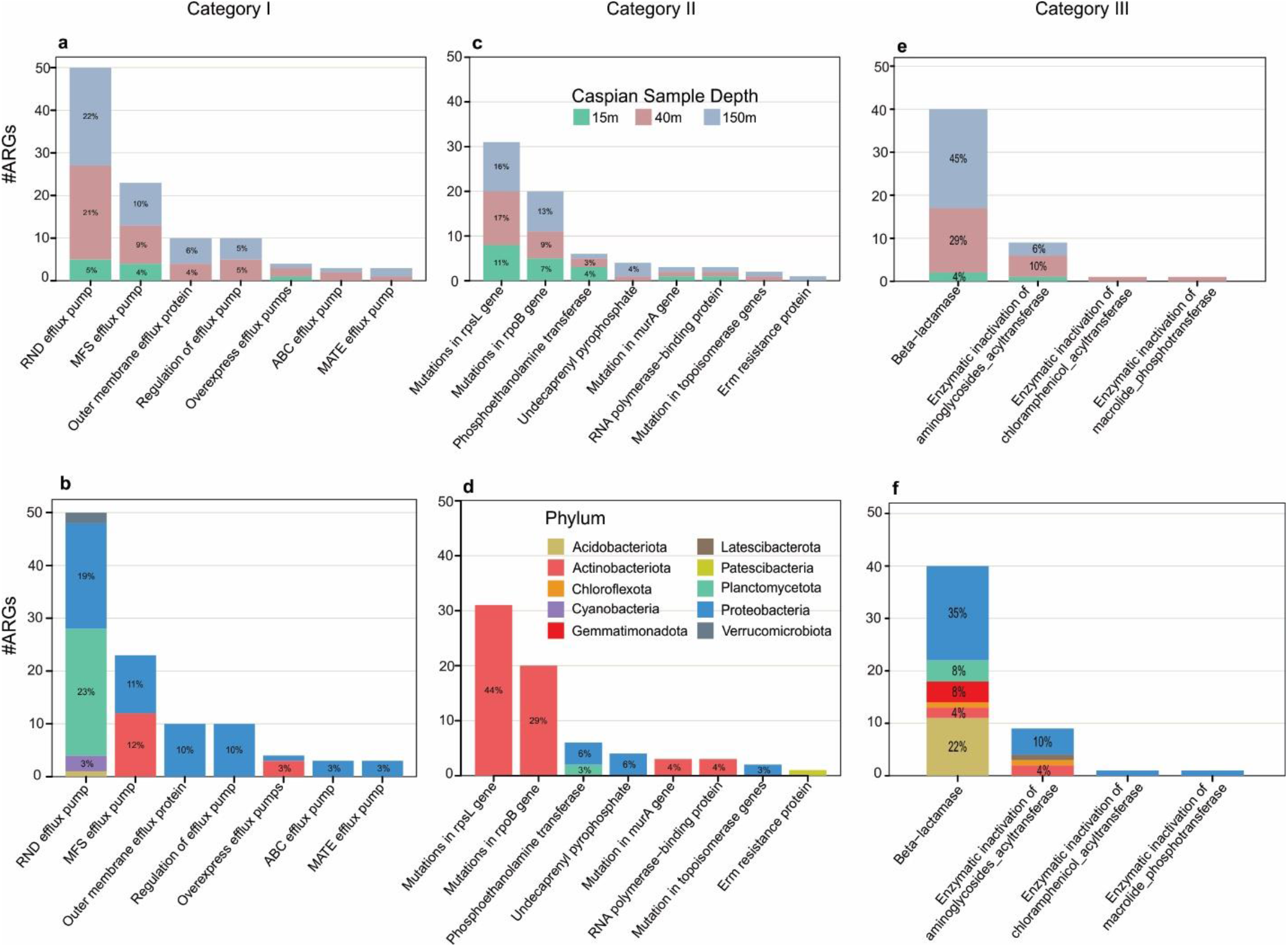
Distribution of ARGs subcategories in Caspian Sea metagenomes based on depth and their phylogenetic diversity at the phylum level. Distribution of ARGs in: **a**,**b** category (I) Prevention of access to target, **c**,**d**, category (II) modification/protection of targets, and **e**,**f**, category (III) direct modification of antibiotics.

We detected six cyclic AMP (cAMP) receptor protein (CRP) genes in the stress resistance category. All of these six stress resistance genes belong to the class Gammaproteobacteria (Fig. 1f) and five of them belong to the family Pseudohongiellaceae. CRP, a global transcriptional regulator, contributes to emergence of stress resistance in bacteria through its regulatory role in multiple cellular pathways, such as anti-oxidation and DNA repair pathways. Stress responses play an important role in integron rearrangements, facilitating the antibiotic resistance acquisition and development, and ultimately the emergence of multidrug-resistant bacteria. So, understanding the evolution of bacterial stress responses is critical, since they have a major impact on the evolution of genome plasticity and antibiotic resistance (27).

A recent study demonstrates the potential contribution of metal resistance genes and plasmidome to the stabilization and persistence of the antibiotic resistome in aquatic environments (28). We identified three ferritin genes classified in the metal resistance category, in the Caspian Sea MAGs affiliated to genus *Mycolicibacterium*. Ferritin (bfr) is an iron storage protein involved in protection of cells against oxidative stress (iron-mediated oxidative toxicity) and iron overload (29,30).

The most frequent subcategory detected in the Caspian Sea MAGs was RND efflux pump (50 ARGs) and after that, β-lactamase (40 ARGs) and mutation in rpsL gene (31 ARGs) (**Table 1**). In category Prevention of access to target, Caspian ARGs are classified into different types of efflux pumps and their regulatory sequences (Fig. 2a, b). Besides, all resistance genes caused by mutational changes are in category modification/protection of targets (Fig. 2c, d). In the category of direct modification of antibiotics, Caspian ARGs belong to β-lactamases and some transferases (Fig. 2e, f). Many ARGs provide resistant to several classes of antibiotics in bacteria, consequently when inferring the antibiotic classes that the Caspian Sea ARGs provide resistance to the highest percentage refer to multidrug antibiotic class (Fig. 3). The genes encoding multidrug efflux pumps are evolutionarily ancient elements and are highly conserved (7). The frequency of the efflux mediated antibiotic resistance in other environments (14,19) supported that efflux pumps have other physiologically relevant roles such as detoxification of intracellular metabolites, stress response and cell homeostasis in the natural ecosystems (7). The second antibiotic class that the Caspian Sea ARGs provide resistance to is the β-lactams class. many soil bacteria have been isolated that can grow on β-lactam antibiotics as the sole source of carbon (31,32). The abundance of β-lactamases in the Caspian Sea MAGs could also be related to other ecological roles of β-lactam.

**Figure 3.**
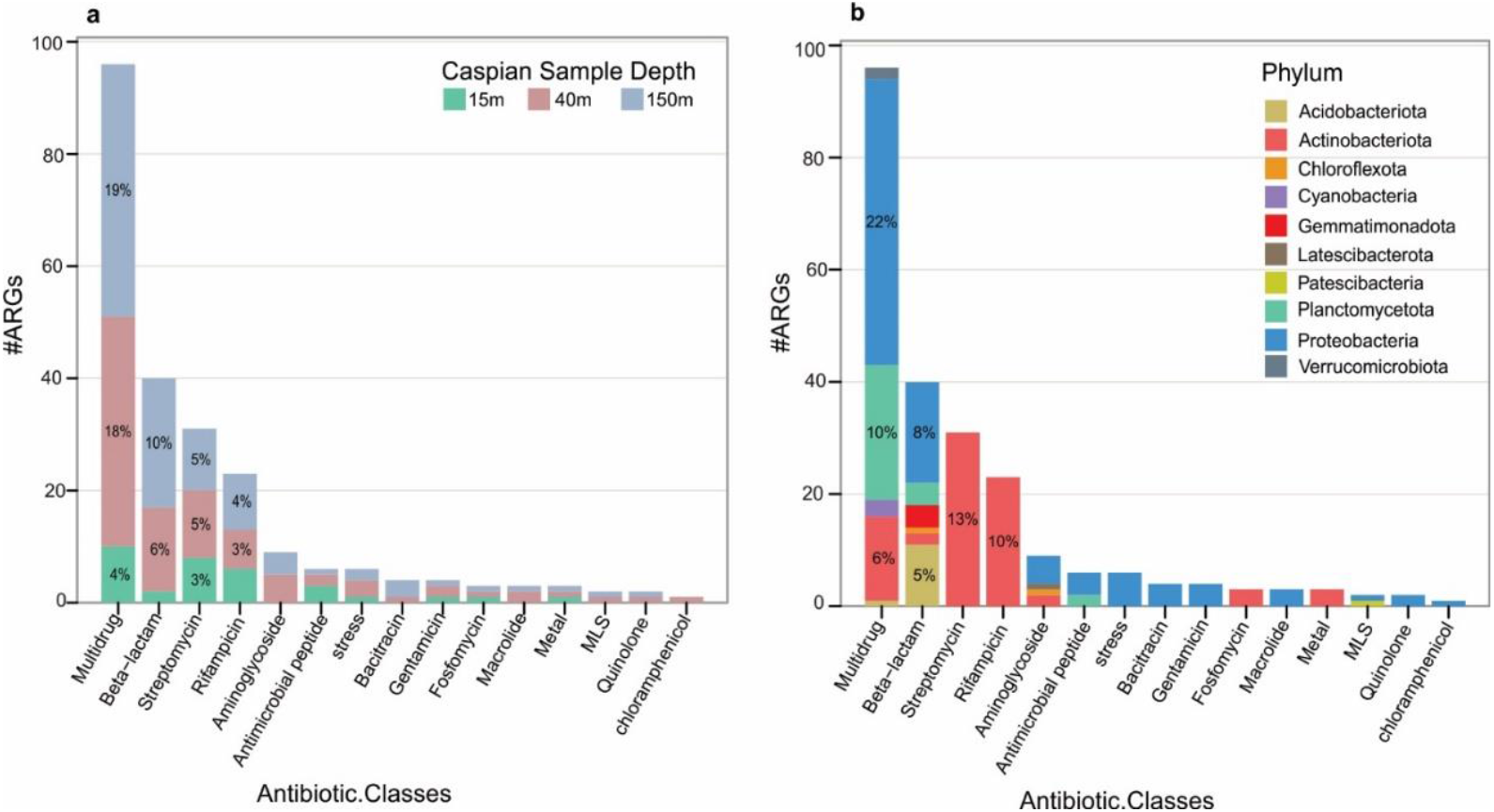
Abundance and distribution of drug classes that the antibiotic resistance genes of the Caspian Sea metagenomes can provide resistance against. Distribution of drug classes based on depth (**a**), and phylogenetic diversity at the phylum level (**b**).

### Taxonomic distribution of ARG containing genomes

A total of 233 resistance genes were identified from 102 reconstructed MAGs of the Caspian Sea (MAG stats are shown in the **Supplementary Table S3**). These MAGs were assigned to 10 phyla dominated by Actinobacteriota (79 ARGs in 39 ARBs) and Proteobacteria (99 ARGs in 28 ARBs) (Fig. 1b, f). Identified ARGs were distributed in 15 classes showing the highest abundance in Acidimicrobiia (24 ARBs), Gammaproteobacteria (22 ARBs) and Actinobacteria (15 ARBs) (Fig. 1c). Although Bacteroidota constitutes 18% of reconstructed Caspian Sea MAGs, no resistance gene was detected in MAGs affiliated to this phylum (**Supplementary Figure S1**).

The casp40-mb.75 and casp150-mb.119 MAGs contained ARGs belonging to four different groups of resistance genes (**Supplementary Figure S4**). Both MAGs have one contain an ARG in the metal resistance group (and no stress resistance gene). These MAGs are taxonomically affiliated to the genus *Mycolicibacterium* and show a higher abundance at 40m and 150m depths (ca. 1300 TPM) (**Supplementary Figure S5**). The genus *Mycolicibacterium* comprise a wide range of environmental and pathogenic bacteria that are potential hosts of ARGs and MGEs. This may contribute to their diversity and evolution or even to their success as opportunistic pathogens (33). Studies conducted in Japan suggest that livestock could acquire *Mycolicibacterium peregrinum* from their environment (34). Presence of *Mycolicibacterium* representatives containing a set of ARGs in the natural environment could be a reservoir of genes for potential development of resistance in pathogenic groups.

Two MAGs affiliated to Pseudomonadales order (casp40-mb.215 and casp150-mb.169) contain ARGs belonging to categories I, II and III (**Supplementary Figure S4**). Among 22 ARG containing MAGs affiliated to Gammaproteobacteria, 14 MAGs belonged to Pseudomonadales order with estimated genome sizes in the range of 2.1 to 5.4 Mbp. Among all ARG containing MAGs, the casp40-mb.215 (n=21 ARGs) and casp150-mb.169 (n=18 ARGs) affiliated with *Acinetobacter venetianus* had the highest number of ARGs (**Supplementary Figure S6**). Representatives of genus *Acinetobacter* are commonly found in soil and water (35). This genus contains *Acinetobacter baumannii* that is a pathogen with known antibiotic resistance complications for infection treatment (36).

Representatives of the Acidimicrobiia class are ubiquitous aquatic microbes with high relative abundances in the brackish Caspian Sea (26). These MAGs have the estimated genome size in the range of 1.3 to 2.9 Mbp and their ARGs belong to the category II and are mainly caused by mutations (Fig. 4 and **Supplementary Figure S3**). In addition to the Acidimicrobiia class, there is a high frequency of antibiotic resistance mechanisms based on target modification and protection detected in the Actinobacteria affiliated MAGs (Fig. 4). For streamlined members of this taxon that are highly abundant in the ecosystem and have adapted to the oligotrophic environments, it could potentially be advantageous to modify the antibiotics target to develop antibiotic resistance so they can avoid the cost of carrying a new gene for developing resistant phenotype. 12 MAGs affiliated to Nanopelagicales order in Actinobacteria class contain 20 ARGs. All these ARGs are in category II and subcategories mutation in rpoB and rpsL genes (12 rpoB gene and 8 rpsL gene). Unlike other Actinobacteria, members of the order Nanopelagicales, family Nanopelagicaceae and AcAMD-5, have a low G + C% content (38% to 47%) in their genome and have streamlined genomes in the range of 1.3 to 1.6 Mbp. Members of this order are present in freshwater and brackish environments such as the Caspian Sea in high abundances making up more than 30% of the microbial community in the surface layer of freshwater ecosystems (37). According to streamlining theory, these organisms remove unnecessary genes from their genomes, thereby lowering the cellular metabolic costs (38). In line with this strategy, the use of antibiotic resistance mechanisms based on modification or protection of the target, especially based on mutations in the antibiotic target, seems to be one of the best options to achieve antibiotic resistance in members of such lineages (**Supplementary Figure S3**). Although this order does not contain a known pathogenic representative, their ubiquitously high abundance in the ecosystem could offer a new perspective on the ecological role of antibiotic resistance genes. Although the family S36-B12 from order Nanopelagicales is one of the high G + C% content (about 60%) groups with the estimated genome size in the range of 2 to 3 Mbp, they also developed their resistance through mutations. While we attribute the resistant streamlined genomes to the target modification and protection mechanisms (especially based on mutations), this may be a feature of class Actinobacteria.

**Figure 4.**
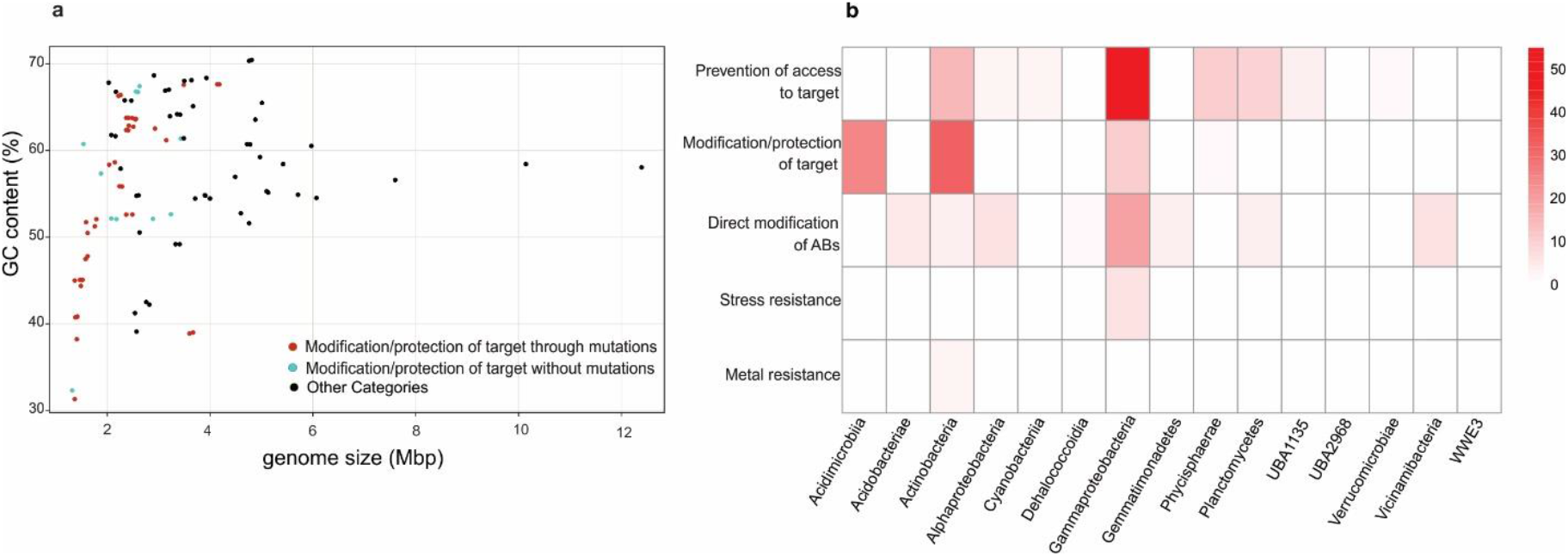
Genome streamlining in ARG containing MAGs. **a**, Genomic GC content versus estimated genome size for all ARG containing MAGs, red dots indicate genomes that confer category II resistant through mutations, which have lower GC content and estimated genome size. Blue dots indicate genomes that confer category II resistant without mutation. Other categories are shown in **Supplementary Figure S3. b**, Heat map representation of number of genomes at the taxonomic level of class and ARG categories. Classes of Acidimicrobiia and Actinobacteria have a higher number of ARG category II.

Among all ARG containing MAGs, class Acidimicrobiia affiliated MAGs show the highest abundance in three depths of Caspian Sea followed by class Actinobacteria (**Supplementary Figure S5**). The MAG of casp15-mb.93 and casp15-mb.71 are among the most abundant bacteria with detected ARGs in 40 and 150 m depth metagenomes. These MAGs belong to order Microtrichales and their ARGs were classified in category II having mutations in the rpsL genes. While these MAGs were reconstructed from the 15m depth metagenomes, they show a higher abundance at the lower strata (**Supplementary Figure S5**).

Prior culture based studies on the Zarjoub (39) and Gowharrood (40) rivers that are entering the Caspian Sea basin report antibiotic resistant coliform bacteria. These studies do not report the antibiotic concentrations of the natural environment but claim that presence of antibiotic resistant bacteria is due to uncontrolled discharge of agricultural and livestock effluents upstream of the river and the entry of municipal and hospital wastewater into these two rivers and later the Caspian Sea. Hence it is important to understand the accurate resistome profile of this natural ecosystem as a step toward sustaining its ecosystem services.

The Caspian Sea ARG containing MAGs are dominated by representatives of Acidimicrobiia, Gammaproteobacteria and Actinobacteria classes. A recent study on the deep-sea water (more than 1000 m deep) suggest that even deep marine environments could be an environmental reservoir for ARGs mainly carried by representatives of Gammaproteobacteria (70%) and Alphaproteobacteria (20%)(19). The identified ARGs were classified based on the classes of antibiotics they provide resistance to and most abundant identified ARG types respectively included multidrug, peptide and aminoglycoside (19). Exploring the diversity and abundance of ARGs in global ocean metagenomes using machine-learning approach (DeepARG tool) showed that ARGs conferring resistance to tetracycline are the most widespread followed by those providing resistance to multidrug and β-lactams. In the contigs containing ARGs, Alphaproteobacteria was identified as the largest taxonomic unit, followed by Gammaproteobacteria (14). In the Caspian Sea however, similar to the global ocean (14) most identified ARGs provide resistance to multidrug class followed by β-lactams (Fig. 3). Caspian ARGs conferring resistance to tetracycline were annotated as transporter groups and consequently we classified them into category I and multidrug class. We explored the distribution of ARGs in Caspian viral contigs using the same method, six viral contigs identified by the virsorter2 contained ARGs however in the follow up manual curations we could not confirm the viral origin of these contigs and removed them from the results.

### Phylogenetic analysis of the Caspian Sea β-lactamases

A total of 40 ARGs classified as β-lactamase genes (*bla*), were detected in the MAGs of the Caspian Sea and they phylogenetic relations were analyzed (reference sequences and tree file are accessible in **Supplementary Data File S1**). Beta-lactamases are classified into four molecular classes based on their amino acid sequences (A to D classes). Class A, C, and D enzymes utilize serine for β-lactam hydrolysis and class B are metalloenzymes that require divalent zinc ions for substrate hydrolysis (41). Among identified Caspian *bla*, 9, 22, 2, and 7 are classified in respectively class A, B, C, and D (**Supplementary Table S4**). As shown in Fig. 5, most of the Caspian *bla* are metallo-β-lactamases. Metallo-beta-lactamase enzymes pose a particular challenge to drug development due to their structure and diversity (42). These enzymes escape most of the recently licensed beta-lactamase inhibitors. Acquired metallo-beta-lactamases, which are prevalent in Enterobacterales and *Pseudomonas aeruginosa*, are usually associated with highly drug-resistant phenotypes and are more dangerous (42). While the *bla* containing reference genes included in this phylogeny (collected from the KEGG database) mostly belong to the Gammaproteobacteria, Caspian *bla* containing genomes represent a higher diverse belonging to six different phyla (in 8 different classes). Some of these *bla* containing MAGs are affiliated to taxa that do not yet have a representative in culture. natural ecosystems are known to be important reservoirs of β-lactamase gene homologs, however, exchange of β-lactamases between natural environments and human and bovine fecal microbiomes occurs at low frequencies (43). Additionally, β-lactams can be used as a source of nutrient after β-lactamase cleavage. The β-lactam catabolism pathway has been detected in diverse Proteobacteria isolates from soil that is generating carbon sources for central metabolism (32,44).

**Figure 5.**
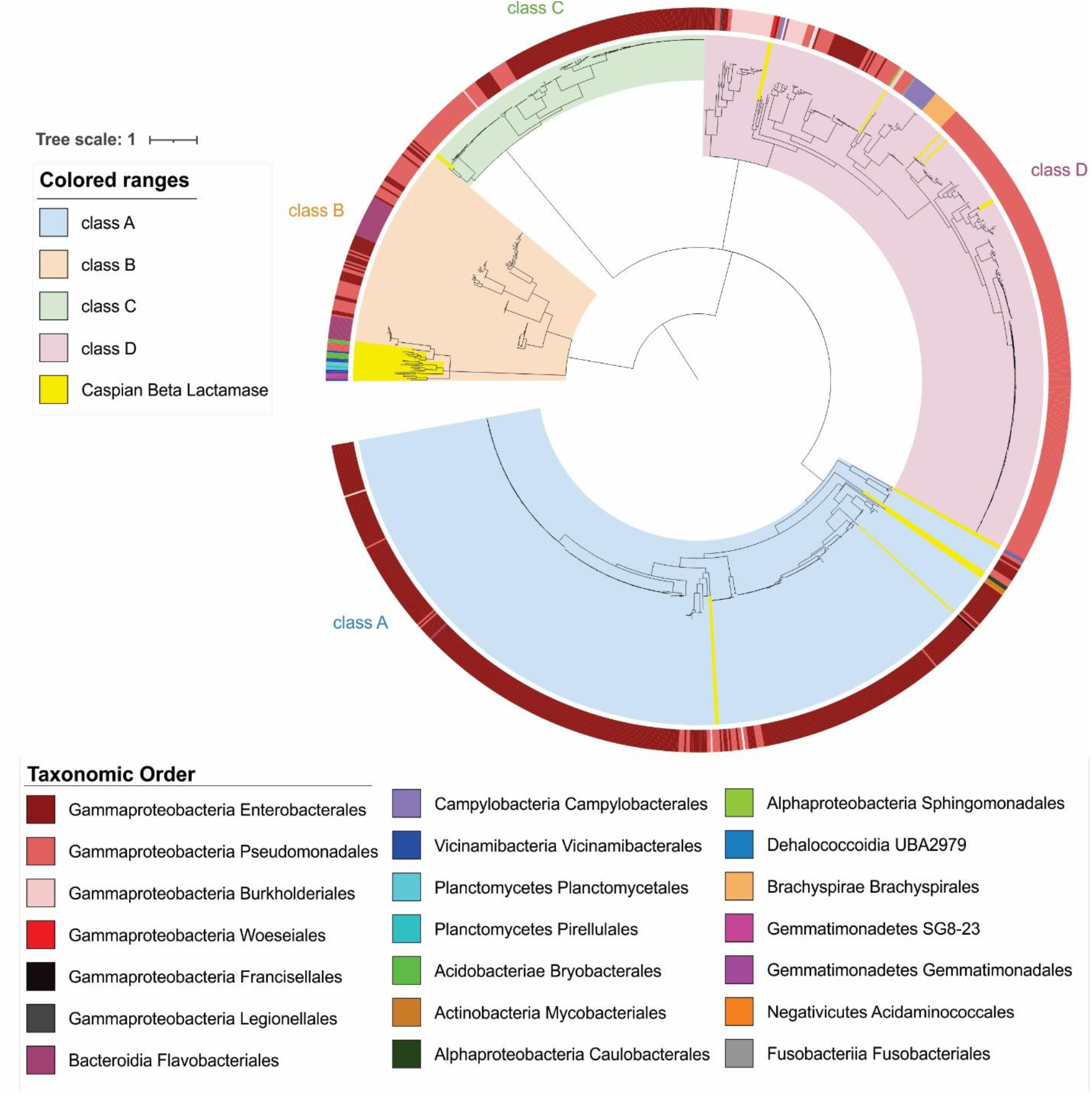
Maximum-likelihood phylogenetic tree of β-lactamases. β-lactamases are classified into four classes based on their amino acid sequences (A to D classes). Phylogenetic tree was constructed by using the maximum likelihood method, and 100 bootstrap replications. Taxonomy of *bla* containing genomes at the order level is annotated on the phylogenetic tree. Caspian β-lactamases are highlight in yellow in the tree.

## Conclusions

Antibiotic resistance is a global health challenge and according to One Health approach, attention to environmental antibiotic resistome is critical to combat AMR. Our study shows the distribution of antibiotic resistance genes in the Caspian Sea ecosystem, even though no accurate measurement of antibiotic contamination of the Caspian Sea has been reported so far. Moreover, our findings revealed the mechanism of resistant streamlined genomes which is based on target modifications. The resistome profile and the type of resistance mechanism of the Caspian Sea MAGs provided in this study can be used as a reference database for monitoring the development and spread of antibiotic resistance in the Caspian Sea over time and can also guide future studies. The increase of antibiotic concentrations in natural ecosystems, as a consequence of human activities, not only influences the Prevalence of antibiotic resistance genes, but also can alter the microbial populations and communities of the Caspian Sea. It can have adverse effects on the carbon and nitrogen cycle balance and hence may lead to imbalances in the homeostasis of microbial communities in the Caspian Sea leading to potentially severe consequences for this ecosystem as a whole. However, as bacterial communities are formed by a complex array of evolutionary, ecological and environmental factors, it is difficult to obtain a clear understanding of the evolution and ecology of antibiotic resistance in natural environments. Eventually, Global problems require global solutions and only a concerted and sustained international effort can succeed in dealing with AMR.

## Methods

### Assembly and binning of the Caspian Sea metagenomes

Brackish Caspian Sea metagenome were used for in-silico screening of ARGs. Metagenomic datasets derived from three different depths of the Caspian Sea (15 m, 40 m, and 150 m), were published in 2016 by Mehrshad et al. (26) and are accessible under the BioProject identifier PRJNA279271. These metagenomes were quality checked using bbduk.sh script (sourceforge.net/projects/bbmap/) and assembled using metaSPAdes (45). Metagenomic reads were mapped against assembled contigs using bbmap.sh script (sourceforge.net/projects/bbmap/). Contigs ≥ 2kb were binned based on differential coverage and composition using Metabat2 (46). Quality of the reconstructed MAGs was assessed using CheckM (47) and bins with completeness ≥ 40% and contamination ≤ 5% were used for further analysis. Taxonomy of these MAGs was assigned using GTDB-tk (v0.3.2) and genome taxonomy database release R89 (48). MAG abundances in different metagenomes of the Caspian sea were calculated using the CoverM tool with transcript per million (TPM) method (https://github.com/wwood/CoverM).

### ARG identification

The ARGs in the Caspian Sea MAGs were determined using the six different pipelines and software (RGI, AMRFinder, ResFinder, sraX, DeepARG, ABRicate equipped with ARG-ANNOT) (**Supplementary Table S1**). Protein coding sequences of each MAG were predicted using Prodigal (49). The protein sequences of the reconstructed MAGs were searched for ARGs against the Comprehensive Antibiotic Resistance Database (CARD) using Web portal RGI 5.1.1, CARD 3.1.1 (https://card.mcmaster.ca/analyze/rgi) with default settings (50).

NCBI AMRFinderPlus v3.9.3 (https://github.com/ncbi/amr/wiki) command line tool and its associated database, The Bacterial Antimicrobial Resistance Reference Gene Database (which contains 4,579 antimicrobial resistance proteins and more than 560 HMMs), were used for screening ARGs. The protein sequences of all reconstructed MAGs were analyzed with parameter “-p” (51). Additionally, all ARGs present in the MAGs protein sequences were screened using a deep learning approach, DeepARG v1.0.2 command line tool, (https://bitbucket.org/gusphdproj/deeparg-ss/src/master/) with DeepARG-DB database (--model LS --type nucl --arg-alignment-identity 60) (52).

The nucleotide sequences of the reconstructed MAGs were searched for ARGs using ResFinder 4.1 command line tool (https://bitbucket.org/genomicepidemiology/resfinder/src/master/) and its associated database, ResFinder database with parameters “-ifa -acq -l 0.6 -t 0.8” (53). They were also searched using ARGminer v1.1.1 database (54) and BacMet v2.0 database (55) using sraX v1.5 command line tool (https://github.com/lgpdevtools/srax) with parameters “-db ext–s blastx” (56). These sequences were also searched against ARG-ANNOT v4 database (57) using ABRicate v0.8 command line tool (58).

Results of these methods presented candidate ARGs in our MAG set. Functions of the ARG candidates were further verified using five different annotation tools (default settings); Batch web conserved domain search (CD-Search) in NCBI https://www.ncbi.nlm.nih.gov/Structure/bwrpsb/bwrpsb.cgi (59), web-based Hmmer v2.41.1 (phmmer) https://www.ebi.ac.uk/Tools/hmmer/search/phmmer (60), hmmscan against Pfam v34.0 database http://pfam.xfam.org/search#tabview=tab1 (61), GhostKOALA v2.2 https://www.kegg.jp/ghostkoala/(62), and eggNOG-mapper v2 http://eggnog-mapper.embl.de/ (63). All functional annotation results were compiled and results were compared to obtain a consensus assignment. Then, ARGs were manually curated into 5 antibiotic resistance categories and 21 subcategories based on their functional annotations. The overall workflow of this study is shown in **Supplementary Figure S7**.

### Gene alignment

To confirm resistance due to mutation events in the candidate Caspian ARGs, multiple amino acid sequence alignment was carried out Using Clustal-W (default parameters) (64) embedded in MEGA-X software (65). For each type of the ARGs, reference gene with specific mutations was downloaded from CARD database (**Supplementary Table S2** shows the detail of mutations involved in antibiotic resistance).

### Beta-lactamase phylogeny

To understand the evolutionary relationship of the recovered β-lactamase enzymes, firstly, 1141 reference protein sequences (beta-Lactamase gene variants) were downloaded from KEGG database, https://www.genome.jp/kegg/annotation/br01553.html and combined with β-lactamases recovered from Caspian MAGs (40 protein sequences). Then, all β-lactamase sequences were subjected to multiple sequence alignment using Clustal-W embedded in MEGA-X (Molecular Evolutionary Genetics Analysis) software (65). Phylogenetic tree was constructed using the maximum likelihood method, JTT matrix-based model, and 100 bootstrap replications in MEGA-X software. The bootstrap consensus tree inferred from 100 replicates is taken to represent the evolutionary history. This analysis involved 1181 amino acid sequences in total. Taxonomic assignment of MAGs was extended to the of β-lactamases and iTOL v6.3.1 was used to annotate and visualize the final phylogenetic tree (66).

### Viral contigs identification

Viral contigs were identified in contigs longer than 1kb using VirSorter2 tool at the score threshold of 0.8 (67). These contigs were further checked manually to ensure the viral origin.

## Data availability

The Caspian Sea metagenomes used for this study have been deposited to GenBank by Mehrshad et al. (26) and are accessible via the bioproject PRJNA279271. Genomes containing ARGs were also deposited to GenBank and are accessible under the accession number Bioproject PRJNA279271. All alignments used for phylogeny reconstruction are accompanying this manuscript as supplementary data.

## Acknowledgements

The computational analysis was performed at the Center for High-Performance Computing, School of Mathematics, Statistics, and Computer Science, University of Tehran.

## Author contributions

MM and SA designed the study. ZG and MM performed the bioinformatics analysis and drafted the manuscript. All authors analyzed and interpreted the data and approved the manuscript.

## Supplementary Tables

**Supplementary Table S1.** Summary of ARG identification tools and databases used in this study.

| Tool | Database | Approach | Input |
| --- | --- | --- | --- |
| **RGI** | CARD | BLASTp | Protein sequence |
| **AMRFinder (NCBI)** | The Bacterial Antimicrobial Resistance Reference Gene Database | BLASTp & HMM scan | Protein sequence |
| **DeepARG** | DeepARG-DB | deep-learning | Protein sequence |
| **ResFinder** | ResFinder | BLASTn | Nucleotide sequence |
| **sraX** | CARD & ARGminer & BacMet | BLASTx | Nucleotide sequence |
| **ABRicate** | ARG-ANNOT | BLASTn | Nucleotide sequence |

**Supplementary Table S2.** Reference genes and the mutation position for each of ARGs that confirm resistance due to mutation events.

| Gene symbol | Gene name | # genes | Reference gene | Mutation position | Ref. |
| --- | --- | --- | --- | --- | --- |
| **rpoB** | DNA-directed RNA polymerase subunit beta | 20 | NP_215181<br>CCP43410.1 | L511R<br>D516G<br>H526T, E250G | https://card.mcmaster.ca/ontology/39867 |
| **rpsL** | 30S ribosomal protein S12 | 43 | WP_003910993 | K43R<br>K88R | https://card.mcmaster.ca/ontology/39979 |
| **rpsA** | 30S ribosomal protein S1 | 3 | WP_016810233<br>CCP44394 | D123A<br>Δ438A<br>A412V | https://card.mcmaster.ca/ontology/42776 |
| **gyrA** | DNA gyrase subunit A (Topoisomerase) | 7 | WP_023644204<br>WP_000116442 | A90V, D94G<br>G79C, S81L | https://card.mcmaster.ca/ontology/39879,<br>https://card.mcmaster.ca/ontology/40507 |
| **parC** | Topoisomerase IV subunit A | 2 | WP_000202252 | S84L, D105E | https://card.mcmaster.ca/ontology/40508 |
| **LpxC** | UDP-3-O-acetyl-N-acetylglucosamine deacetylase | 4 | AJF83452 | P30L | https://card.mcmaster.ca/ontology/40184 |
| **murA** | UDP-N-acetylglucosamine enolpyruvyl transferase | 3 | CCE36834 | C117D | https://card.mcmaster.ca/ontology/40467 |

**Supplementary Table S3.** Taxonomy of the Caspian Sea antibiotic resistant MAGs. The percent of completeness and contamination of these MAGs with their estimated genome size, GC content and taxonomic assignments are listed.

| Genome id | Estimated<br>Genome<br>size (Mbp) | GC<br>content<br>% | Completeness<br>% | Contamination<br>% | Taxonomy |
| --- | --- | --- | --- | --- | --- |
| casp150-mb.136 | 5.97 | 60.52 | 92.52 | 3.42 | p__Acidobacteriota;c__Acidobacteriae;o__Bryobacterales;<br>f__UBA6623;g__s__ |
| casp150-mb.154 | 4.72 | 60.73 | 95.67 | 1.83 | p__Acidobacteriota;c__Acidobacteriae;o__Bryobacterales;<br>f__UBA6623;g__s__ |
| casp40-mb.95 | 4.78 | 60.70 | 95.25 | 3.89 | p__Acidobacteriota;c__Acidobacteriae;o__Bryobacterales;<br>f__UBA6623;g__s__ |
| casp150-mb.205 | 3.67 | 65.10 | 93.68 | 4.27 | p__Acidobacteriota;c__Vicimibacteria;o__Vicimibacterales;<br>f__UBA2999;g__s__ |
| casp150-mb.299 | 5.01 | 65.49 | 90.6 | 3.42 | p__Acidobacteriota;c__Vicimibacteria;o__Vicimibacterales;<br>f__UBA8438;g__UBA8438;s__ |
| casp15-mb.71 | 1.76 | 51.25 | 87.09 | 2.56 | p__Actinobacteriota;c__Acidimicrobiia;o__Microtrichales;<br>f__Ilumatobacteraceae;g__BACL27;s__ |
| casp15-mb.93 | 1.59 | 51.73 | 90.62 | 3.68 | p__Actinobacteriota;c__Acidimicrobiia;o__Microtrichales;<br>f__Ilumatobacteraceae;g__BACL27;s__ |
| casp150-mb.152 | 1.62 | 47.79 | 89.32 | 0.85 | p__Actinobacteriota;c__Acidimicrobiia;o__Microtrichales;<br>f__Ilumatobacteraceae;g__BACL27;s__ |
| casp150-mb.247 | 1.62 | 50.47 | 78.66 | 3.42 | p__Actinobacteriota;c__Acidimicrobiia;o__Microtrichales;<br>f__Ilumatobacteraceae;g__BACL27;s__ |
| casp15-mb.34 | 2.42 | 63.76 | 93.16 | 1.28 | p__Actinobacteriota;c__Acidimicrobiia;o__Microtrichales;<br>f__Ilumatobacteraceae;g__Casp-actino5;s__ |
| casp15-mb.84 | 2.49 | 63.73 | 98.29 | 3.85 | p__Actinobacteriota;c__Acidimicrobiia;o__Microtrichales;<br>f__Ilumatobacteraceae;g__Casp-actino5;s__ |
| casp150-mb.11 | 2.55 | 63.57 | 97.44 | 3.94 | p__Actinobacteriota;c__Acidimicrobiia;o__Microtrichales;<br>f__Ilumatobacteraceae;g__Casp-actino5;s__ |
| casp150-mb.170 | 2.37 | 63.76 | 93.16 | 2.14 | p__Actinobacteriota;c__Acidimicrobiia;o__Microtrichales;<br>f__Ilumatobacteraceae;g__Casp-actino5;s__ |
| casp150-mb.321 | 2.22 | 66.30 | 91.45 | 3.85 | p__Actinobacteriota;c__Acidimicrobiia;o__Microtrichales;<br>f__Ilumatobacteraceae;g__Casp-actino5;s__ |
| casp40-mb.212 | 2.56 | 63.64 | 96.58 | 1.28 | p__Actinobacteriota;c__Acidimicrobiia;o__Microtrichales;<br>f__Ilumatobacteraceae;g__Casp-actino5;s__ |
| **casp40-mb.94** | 2.26 | 66.38 | 82.34 | 3.85 | p__Actinobacteriota;c__Acidimicrobiia;o__Microtrichales;<br>f__Ilumatobacteraceae;g__Casp-actino5;s__ |
| **casp15-mb.191** | 2.29 | 55.83 | 94.02 | 1.28 | p__Actinobacteriota;c__Acidimicrobiia;o__Microtrichales;<br>f__Ilumatobacteraceae;g__Casp-actino5;s__Casp-actino5 sp001510385 |
| **casp40-mb.42** | 2.24 | 55.86 | 94.02 | 1.28 | p__Actinobacteriota;c__Acidimicrobiia;o__Microtrichales;<br>f__Ilumatobacteraceae;g__Casp-actino5;s__Casp-actino5 sp001510385 |
| **casp15-mb.86** | 2.41 | 62.32 | 83.59 | 0.85 | p__Actinobacteriota;c__Acidimicrobiia;o__Microtrichales;<br>f__Ilumatobacteraceae;g__Casp-actino8;s__ |
| **casp150-mb.228** | 2.37 | 62.35 | 80.6 | 0.85 | p__Actinobacteriota;c__Acidimicrobiia;o__Microtrichales;<br>f__Ilumatobacteraceae;g__Casp-actino8;s__ |
| **casp40-mb.149** | 2.39 | 62.35 | 83.16 | 0.85 | p__Actinobacteriota;c__Acidimicrobiia;o__Microtrichales;<br>f__Ilumatobacteraceae;g__Casp-actino8;s__ |
| **casp150-mb.258** | 2.37 | 52.60 | 97.61 | 3.07 | p__Actinobacteriota;c__Acidimicrobiia;o__Microtrichales;<br>f__Ilumatobacteraceae;g__Casp-actino8;s__Casp-actino8 sp001510335 |
| **casp40-mb.115** | 2.49 | 52.61 | 96.3 | 2.14 | p__Actinobacteriota;c__Acidimicrobiia;o__Microtrichales;<br>f__Ilumatobacteraceae;g__Casp-actino8;s__Casp-actino8 sp001510335 |
| **casp150-mb.41** | 2.51 | 62.73 | 94.02 | 2.99 | p__Actinobacteriota;c__Acidimicrobiia;o__Microtrichales;<br>f__Ilumatobacteraceae;g__Casp-actino8;s__Casp-actino8 sp001510455 |
| **casp40-mb.285** | 2.42 | 62.86 | 94.02 | 1.71 | p__Actinobacteriota;c__Acidimicrobiia;o__Microtrichales;<br>f__Ilumatobacteraceae;g__Casp-actino8;s__Casp-actino8 sp001510455 |
| **casp40-mb.234** | 2.93 | 62.53 | 93.16 | 2.56 | p__Actinobacteriota;c__Acidimicrobiia;o__Microtrichales;<br>f__Ilumatobacteraceae;g__Ilumatobacter;s__ |
| **casp15-mb.62** | 1.79 | 52.06 | 88.08 | 0.95 | p__Actinobacteriota;c__Acidimicrobiia;o__Microtrichales;<br>f__Ilumatobacteraceae;g__UBA3006;s__UBA3006 sp001438985 |
| **casp150-mb.29** | 1.37 | 31.31 | 75.21 | 2.14 | p__Actinobacteriota;c__Acidimicrobiia;o__TMED189;<br>f__TMED189;g__TMED189;s__ |
| **casp150-mb.86** | 2.04 | 58.36 | 95.73 | 3.56 | p__Actinobacteriota;c__Acidimicrobiia;o__UBA5794;<br>f__UBA5794;g__UBA5794;s__ |
| **casp15-mb.16** | 3.49 | 67.58 | 70.15 | 0.05 | p__Actinobacteriota;c__Actinobacteria;o__Mycobacteriales;<br>f__Mycobacteriaceae;g__Mycolicibacterium;s__ |
| **casp150-mb.119** | 4.18 | 67.64 | 96.53 | 1.61 | p__Actinobacteriota;c__Actinobacteria;o__Mycobacteriales;<br>f__Mycobacteriaceae;g__Mycolicibacterium;s__Mycolicibacterium<br>sp001510415 |
| **casp40-mb.75** | 4.14 | 67.64 | 97.15 | 0.94 | p__Actinobacteriota;c__Actinobacteria;o__Mycobacteriales;<br>f__Mycobacteriaceae;g__Mycolicibacterium;s__Mycolicibacterium<br>sp001510415 |
| **casp40-mb.59** | 1.37 | 44.99 | 67.91 | 1.94 | p__Actinobacteriota;c__Actinobacteria;o__Nanopelagicales; f__AcAMD-5;g__AcAMD-5;s__ |
| **casp150-mb.328** | 1.41 | 38.22 | 64.81 | 0.59 | p__Actinobacteriota;c__Actinobacteria;o__Nanopelagicales; f__AcAMD-5;g__ATZT02;s__ |
| **casp15-mb.194** | 1.51 | 45.04 | 89.92 | 1.23 | p__Actinobacteriota;c__Actinobacteria;o__Nanopelagicales; f__Nanopelagicales;g__AAA044-D11;s__AAA044-D11 sp002340925 |
| **casp150-mb.71** | 1.47 | 45.06 | 86.95 | 0.66 | p__Actinobacteriota;c__Actinobacteria;o__Nanopelagicales; f__Nanopelagicales;g__AAA044-D11;s__AAA044-D11 sp002340925 |
| **casp40-mb.60** | 1.52 | 45.07 | 90.19 | 1.89 | p__Actinobacteriota;c__Actinobacteria;o__Nanopelagicales; f__Nanopelagicales;g__AAA044-D11;s__AAA044-D11 sp002340925 |
| **casp40-mb.83** | 1.49 | 44.37 | 83.65 | 0.86 | p__Actinobacteriota;c__Actinobacteria;o__Nanopelagicales; f__Nanopelagicales;g__MAG-120802;s__ |
| **casp15-mb.46** | 1.42 | 40.84 | 71.59 | 0.79 | p__Actinobacteriota;c__Actinobacteria;o__Nanopelagicales; f__Nanopelagicales;g__Nanopelagicus;s__Nanopelagicus sp001437855 |
| **casp150-mb.270** | 1.38 | 40.74 | 66.2 | 2.81 | p__Actinobacteriota;c__Actinobacteria;o__Nanopelagicales; f__Nanopelagicales;g__Nanopelagicus;s__Nanopelagicus sp001437855 |
| **casp40-mb.272** | 1.41 | 40.76 | 74.4 | 0.7 | p__Actinobacteriota;c__Actinobacteria;o__Nanopelagicales; f__Nanopelagicales;g__Nanopelagicus;s__Nanopelagicus sp001437855 |
| **casp150-mb.195** | 1.58 | 47.47 | 88.52 | 0.7 | p__Actinobacteriota;c__Actinobacteria;o__Nanopelagicales; f__Nanopelagicales;g__Planktophilia;s__ |
| **casp40-mb.119** | 2.15 | 58.63 | 94.61 | 0.81 | p__Actinobacteriota;c__Actinobacteria;o__Nanopelagicales; f__S36-B12;g__GCA-2737125;s__ |
| **casp15-mb.155** | 3.15 | 61.17 | 76.47 | 2.34 | p__Actinobacteriota;c__Actinobacteria;o__Nanopelagicales; f__S36-B12;g__S36-B12;s__ |
| **casp150-mb.221** | 2.26 | 57.90 | 82.84 | 2.42 | p__Chloroflexota;c__Dehalococcoidia;o__SAR202; f__;g__;s__ |
| **casp150-mb.66** | 2.03 | 67.82 | 51.91 | 4.13 | p__Chloroflexota;c__Dehalococcoidia;o__UBA2979; f__UBA2979;g__;s__ |
| **casp15-mb.81** | 5.10 | 55.29 | 63.68 | 3.12 | p__Cyanobacteria;c__Cyanobacteriia;o__Phormidiales; f__Phormidiales;g__;s__ |
| **casp40-mb.69** | 5.13 | 55.14 | 87.27 | 1.63 | p__Cyanobacteria;c__Cyanobacteriia;o__Phormidiales; f__Phormidiales;g__;s__ |
| **casp40-mb.232** | 4.97 | 59.24 | 57.97 | 0.72 | p__Cyanobacteria;c__Cyanobacteriia;o__Phormidiales; f__Phormidiales;g__PCC-6406;s__ |
| **casp150-mb.31** | 3.22 | 63.95 | 97.8 | 2.2 | p__Gemmatimonadota;c__Gemmatimonadetes;o__Gemmatimonadales; f__Gemmatimonadaceae;g__Fen-1231;s__ |
| **casp150-mb.298** | 3.42 | 64.13 | 97.8 | 4.4 | p__Gemmatimonadota;c__Gemmatimonadetes;o__SG8-23; f__UBA6960;g__;s__ |
| **casp40-mb.311** | 3.36 | 64.19 | 98.9 | 4.95 | p__Gemmatimonadota;c__Gemmatimonadetes;o__SG8-23; f__UBA6960;g__;s__ |
| **casp40-mb.18** | 5.71 | 54.89 | 79.83 | 1.2 | p__Latescibacterota;c__UBA2968;o__UBA2968; f__GCA-2709665;g__;s__ |
| **casp150-mb.130** | 1.32 | 32.31 | 55.12 | 0.99 | p__Patescibacteria;c__WWE3;o__f__g__s__ |
| **casp15-mb.10** | 2.91 | 68.66 | 50.24 | 3.98 | p__Planctomycetota;c__Phycisphaerae;o__Phycisphaerales;<br>f__SM1A02;g__s__ |
| **casp150-mb.166** | 2.63 | 50.54 | 96.15 | 0 | p__Planctomycetota;c__Phycisphaerae;o__Phycisphaerales;<br>f__SM1A02;g__s__ |
| **casp150-mb.294** | 2.47 | 65.75 | 88.35 | 1.14 | p__Planctomycetota;c__Phycisphaerae;o__Phycisphaerales;<br>f__SM1A02;g__s__ |
| **casp150-mb.65** | 1.88 | 57.34 | 95.45 | 0 | p__Planctomycetota;c__Phycisphaerae;o__Phycisphaerales;<br>f__SM1A02;g__s__ |
| **casp40-mb.123** | 2.34 | 65.79 | 96.59 | 1.14 | p__Planctomycetota;c__Phycisphaerae;o__Phycisphaerales;<br>f__SM1A02;g__s__ |
| **casp150-mb.61** | 3.49 | 61.40 | 86.53 | 1.24 | p__Planctomycetota;c__Phycisphaerae;o__Phycisphaerales;<br>f__SM1A02;g__UBA12014;s__ |
| **casp40-mb.35** | 3.43 | 61.37 | 93.62 | 2.01 | p__Planctomycetota;c__Phycisphaerae;o__Phycisphaerales;<br>f__SM1A02;g__UBA12014;s__ |
| **casp150-mb.46** | 10.14 | 58.42 | 96.48 | 3.45 | p__Planctomycetota;c__Planctomycetes;o__Pirellulales;<br>f__Pirellulaceae;g__GCA-2726245;s__ |
| **casp150-mb.201** | 3.50 | 68.03 | 90.74 | 1.72 | p__Planctomycetota;c__Planctomycetes;o__Pirellulales;<br>f__UBA1268;g__s__ |
| **casp150-mb.45** | 4.88 | 63.56 | 70.35 | 2.4 | p__Planctomycetota;c__Planctomycetes;o__Pirellulales;<br>f__UBA1268;g__s__ |
| **casp40-mb.39** | 3.64 | 68.10 | 97.63 | 1.15 | p__Planctomycetota;c__Planctomycetes;o__Pirellulales;<br>f__UBA1268;g__s__ |
| **casp150-mb.69** | 12.39 | 58.04 | 62.5 | 1.79 | p__Planctomycetota;c__Planctomycetes;o__Planctomycetales;<br>f__Planctomycetaceae;g__s__ |
| **casp40-mb.109** | 6.07 | 54.52 | 97.69 | 1.19 | p__Planctomycetota;c__Planctomycetes;o__Planctomycetales;<br>f__Planctomycetaceae;g__s__ |
| **casp40-mb.51** | 7.60 | 56.59 | 64.69 | 1.25 | p__Planctomycetota;c__Planctomycetes;o__Planctomycetales;<br>f__Planctomycetaceae;g__s__ |
| **casp150-mb.290** | 2.57 | 54.76 | 95.16 | 1.08 | p__Planctomycetota;c__UBA1135;o__UBA1135;f__UBA1135;g__GCA-2746235;s__ |
| **casp40-mb.240** | 2.61 | 54.82 | 94.09 | 1.08 | p__Planctomycetota;c__UBA1135;o__UBA1135;f__UBA1135;g__GCA-2746235;s__ |
| **casp15-mb.161** | 4.81 | 70.44 | 95.45 | 1.14 | p__Planctomycetota;c__UBA1135;o__UBA2386;f__UBA2386;g__s__ |
| **casp40-mb.29** | 4.76 | 70.34 | 89.2 | 0 | p__Planctomycetota;c__UBA1135;o__UBA2386;f__UBA2386;g__s__ |
| **casp150-mb.133** | 3.20 | 67.01 | 71.35 | 3.01 | p__Proteobacteria;c__Alphaproteobacteria;o__Caulobacterales;<br>f__Caulobacteraceae;g__Brevundimonas;s__ |
| **casp40-mb.174** | 3.13 | 66.89 | 85.68 | 1.1 | p__Proteobacteria;c__Alphaproteobacteria;o__Caulobacterales;<br>f__Caulobacteraceae;g__Brevundimonas;s__ |
| **casp40-mb.87** | 4.49 | 56.96 | 71.02 | 3.75 | p__Proteobacteria;c__Alphaproteobacteria;o__Rhodospirillales_A;<br>f__g__s__ |
| **casp40-mb.99** | 3.93 | 68.36 | 97.85 | 1.08 | p__Proteobacteria;c__Alphaproteobacteria;o__SP197; f__SP197;g__s__ |
| **casp40-mb.200** | 2.54 | 41.23 | 43.86 | 0.43 | p__Proteobacteria;c__Alphaproteobacteria;o__Sphingomonadales;<br>f__Emcibacteraceae;g__UBA4441;s__ |
| **casp150-mb.269** | 2.17 | 66.77 | 75.46 | 1.2 | p__Proteobacteria;c__Alphaproteobacteria;o__UBA2966; f__g__s__ |
| **casp150-mb.97** | 2.60 | 66.74 | 98.1 | 1.92 | p__Proteobacteria;c__Gammaproteobacteria;o__Burkholderiales;<br>f__SG8-39;g__RBG-16-66-20;s__ |
| **casp40-mb.55** | 2.56 | 66.81 | 80.6 | 4.72 | p__Proteobacteria;c__Gammaproteobacteria;o__Burkholderiales;<br>f__SG8-39;g__RBG-16-66-20;s__ |
| **casp150-mb.292** | 2.63 | 67.41 | 88.63 | 1.53 | p__Proteobacteria;c__Gammaproteobacteria;o__Burkholderiales;<br>f__SG8-39;g__SG8-39;s__ |
| **casp15-mb.175** | 2.57 | 39.10 | 91.2 | 0.29 | p__Proteobacteria;c__Gammaproteobacteria;o__Francisellales;<br>f__Francisellaceae;g__Caedibacter;s__ |
| **casp40-mb.114** | 4.60 | 52.76 | 94.8 | 4.28 | p__Proteobacteria;c__Gammaproteobacteria;o__Ga0077536;<br>f__Ga0077536;g__UBA11873;s__ |
| **casp150-mb.297** | 1.54 | 60.75 | 92.49 | 1.17 | p__Proteobacteria;c__Gammaproteobacteria;o__Ga0077554;<br>f__Ga007554;g__s__ |
| **casp40-mb.214** | 2.18 | 52.07 | 81 | 0.66 | p__Proteobacteria;c__Gammaproteobacteria;o__Pseudomonadales;<br>f__Halieaceae;g__IMCC3088;s__ |
| **casp15-mb.124** | 2.08 | 52.15 | 71.43 | 1.49 | p__Proteobacteria;c__Gammaproteobacteria;o__Pseudomonadales;<br>f__Halieaceae;g__IMCC3088;s__IMCC3088 sp003520285 |
| **casp15-mb.181** | 3.24 | 52.64 | 79.8 | 3.33 | p__Proteobacteria;c__Gammaproteobacteria;o__Pseudomonadales;<br>f__Halieaceae;g__Luminiphilus;s__ |
| **casp40-mb.140** | 5.42 | 58.42 | 87.98 | 4.69 | p__Proteobacteria;c__Gammaproteobacteria;o__Pseudomonadales;<br>f__HTCC2089;g__s__ |
| **casp150-mb.241** | 2.82 | 42.21 | 75.66 | 1.4 | p__Proteobacteria;c__Gammaproteobacteria;o__Pseudomonadales;<br>f__Moraxellaceae;g__Acinetobacter;s__Acinetobacter johnsonii |
| **casp40-mb.142** | 2.76 | 42.52 | 59.01 | 0.83 | p__Proteobacteria;c__Gammaproteobacteria;o__Pseudomonadales;<br>f__Moraxellaceae;g__Acinetobacter;s__Acinetobacter johnsonii |
| **casp150-mb.169** | 3.60 | 38.87 | 98.18 | 1.73 | p__Proteobacteria;c__Gammaproteobacteria;o__Pseudomonadales;<br>f__Moraxellaceae;g__Acinetobacter;s__Acinetobacter venetianus |
| **casp40-mb.215** | 3.67 | 38.99 | 98.53 | 0.79 | p__Proteobacteria;c__Gammaproteobacteria;o__Pseudomonadales;<br>f__Moraxellaceae;g__Acinetobacter;s__Acinetobacter venetianus |
| **casp15-mb.5** | 2.89 | 52.11 | 94.81 | 2.16 | p__Proteobacteria;c__Gammaproteobacteria;o__Pseudomonadales;<br>f__Pseudohongiellaceae;g__OM182;s__ |
| **casp15-mb.167** | 3.91 | 54.77 | 95 | 1.17 | p__Proteobacteria;c__Gammaproteobacteria;o__Pseudomonadales;<br>f__Pseudohongiellaceae;g__UBA9145;s__ |
| **casp150-mb.100** | 4.76 | 51.60 | 94.63 | 1.98 | p__Proteobacteria;c__Gammaproteobacteria;o__Pseudomonadales;<br>f__Pseudohongiellaceae;g__UBA9145;s__ |
| **casp150-mb.148** | 3.33 | 49.18 | 94.07 | 1.79 | p__Proteobacteria;c__Gammaproteobacteria;o__Pseudomonadales;<br>f__Pseudohongiellaceae;g__UBA9145;s__ |
| **casp40-mb.220** | 3.90 | 54.83 | 90.06 | 1.94 | p__Proteobacteria;c__Gammaproteobacteria;o__Pseudomonadales;<br>f__Pseudohongiellaceae;g__UBA9145;s__ |
| **casp40-mb.61** | 3.41 | 49.18 | 94.07 | 1.23 | p__Proteobacteria;c__Gammaproteobacteria;o__Pseudomonadales;<br>f__Pseudohongiellaceae;g__UBA9145;s__ |
| **casp150-mb.9** | 4.00 | 54.45 | 89.06 | 2.81 | p__Proteobacteria;c__Gammaproteobacteria;o__Woeseiales;<br>f__Woeseiaceae;g__SZUA-117;s__ |
| **casp40-mb.225** | 3.71 | 54.45 | 88.41 | 2.51 | p__Proteobacteria;c__Gammaproteobacteria;o__Woeseiales;<br>f__Woeseiaceae;g__SZUA-117;s__ |
| **casp150-mb.222** | 2.16 | 61.65 | 97.97 | 0.68 | p__Verrucomicrobiota;c__Verrucomicrobiae;o__Chthoniobacterales;<br>f__;g__;s__ |
| **casp40-mb.44** | 2.08 | 61.78 | 97.8 | 1.01 | p__Verrucomicrobiota;c__Verrucomicrobiae;o__Chthoniobacterales;<br>f__;g__;s__ |

**Supplementary Table S4.**
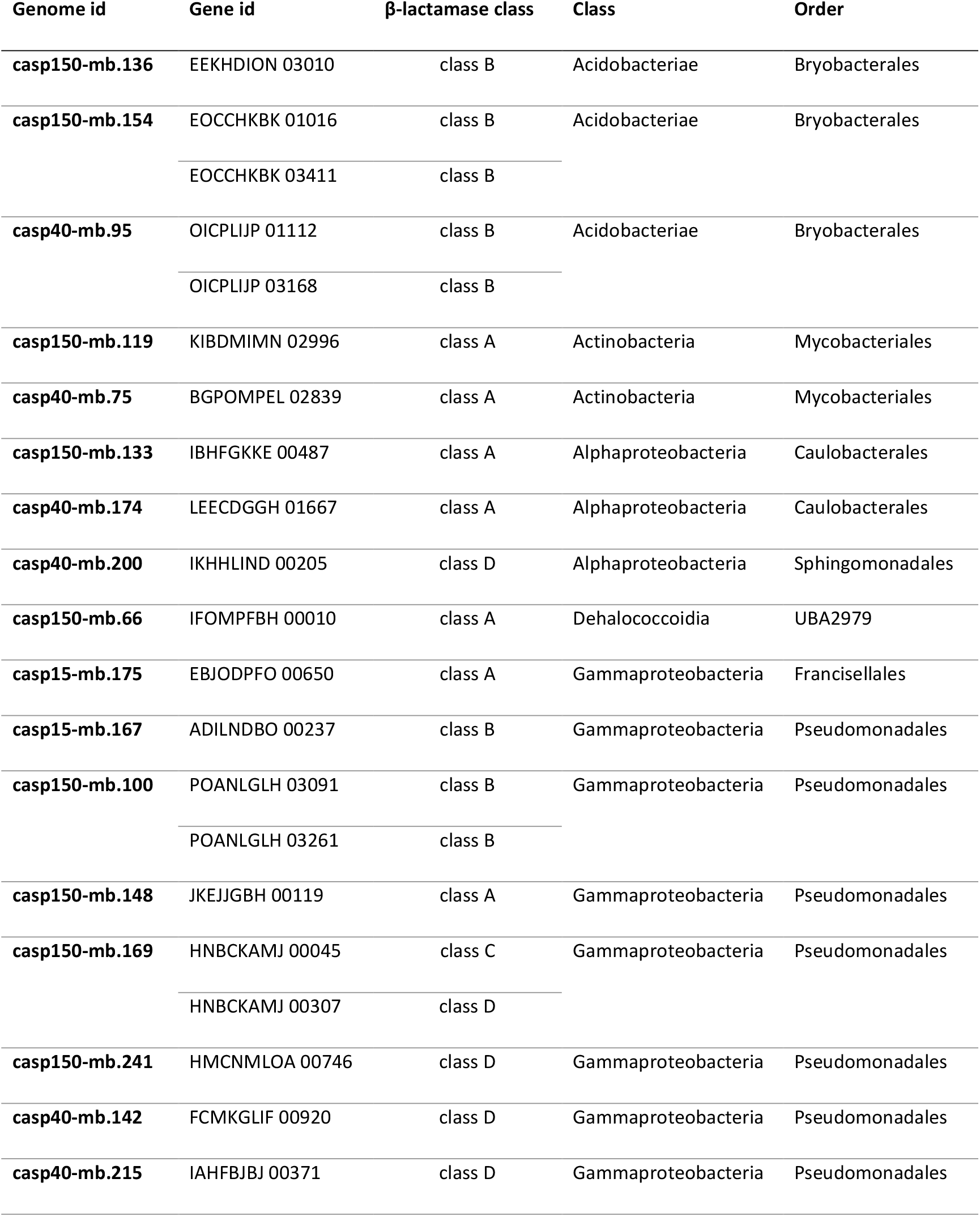
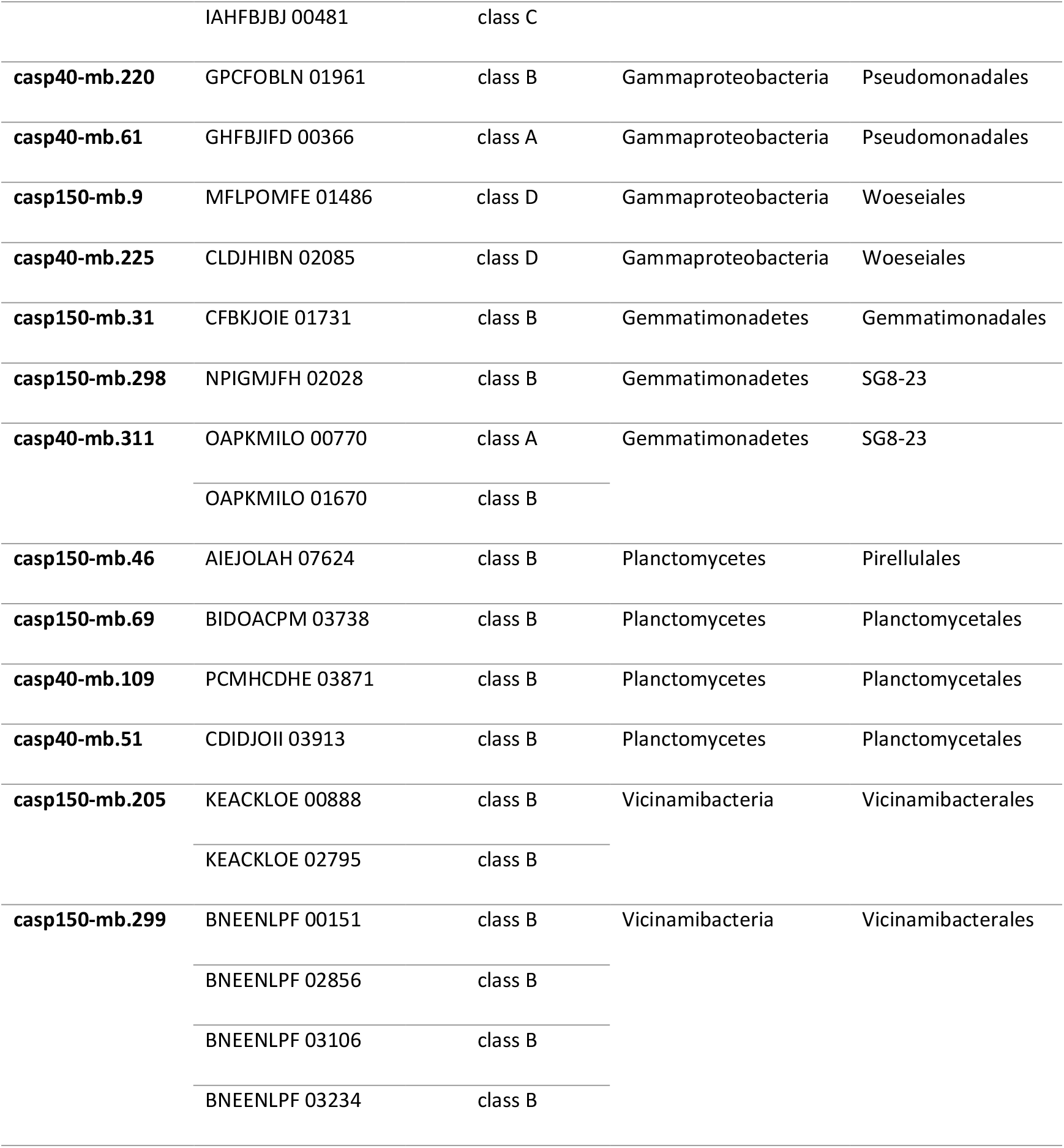
List of β-lactamase containing genomes in the Caspian Sea MAGs and their taxonomic affiliation. There are 40 β-lactamase genes detected in 30 Caspian Sea MAGs.

## Supplementary figures

**Figure S1.**
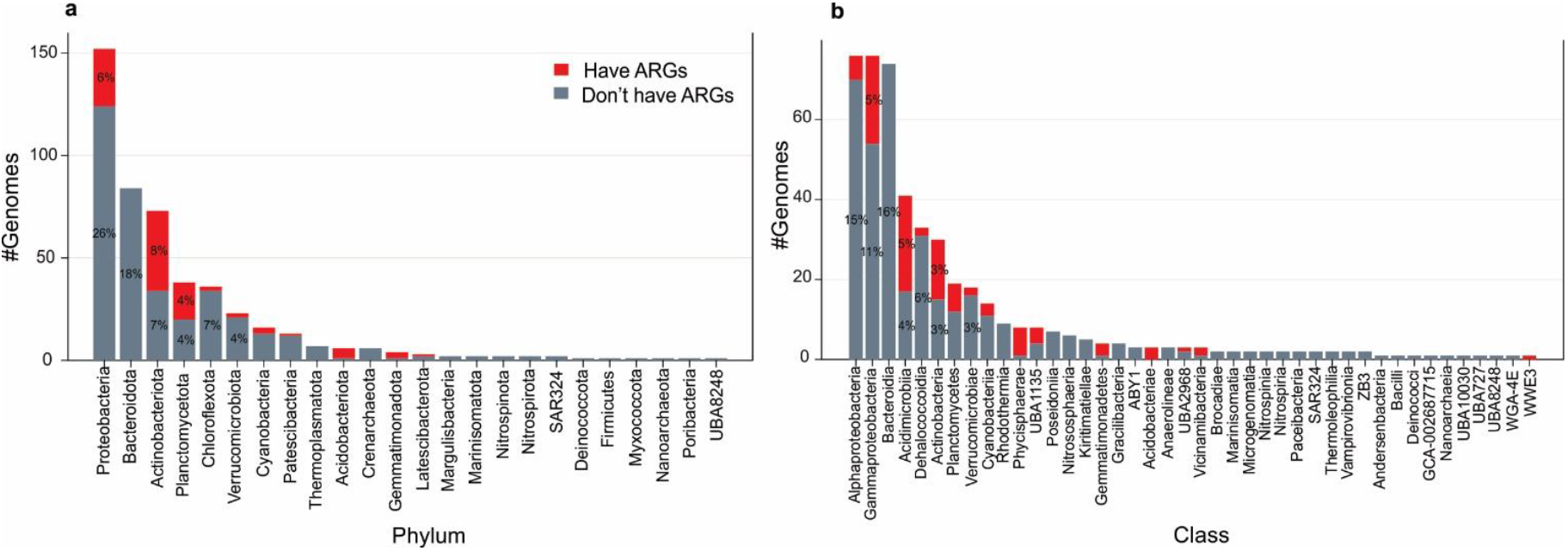
Overall taxonomic distribution of all reconstructed MAGs from the Caspian Sea metagenomes at phylum (**a**) and class (**b**) level. The red areas indicate the number of ARG containing MAGs in each taxa.

**Figure S2.**
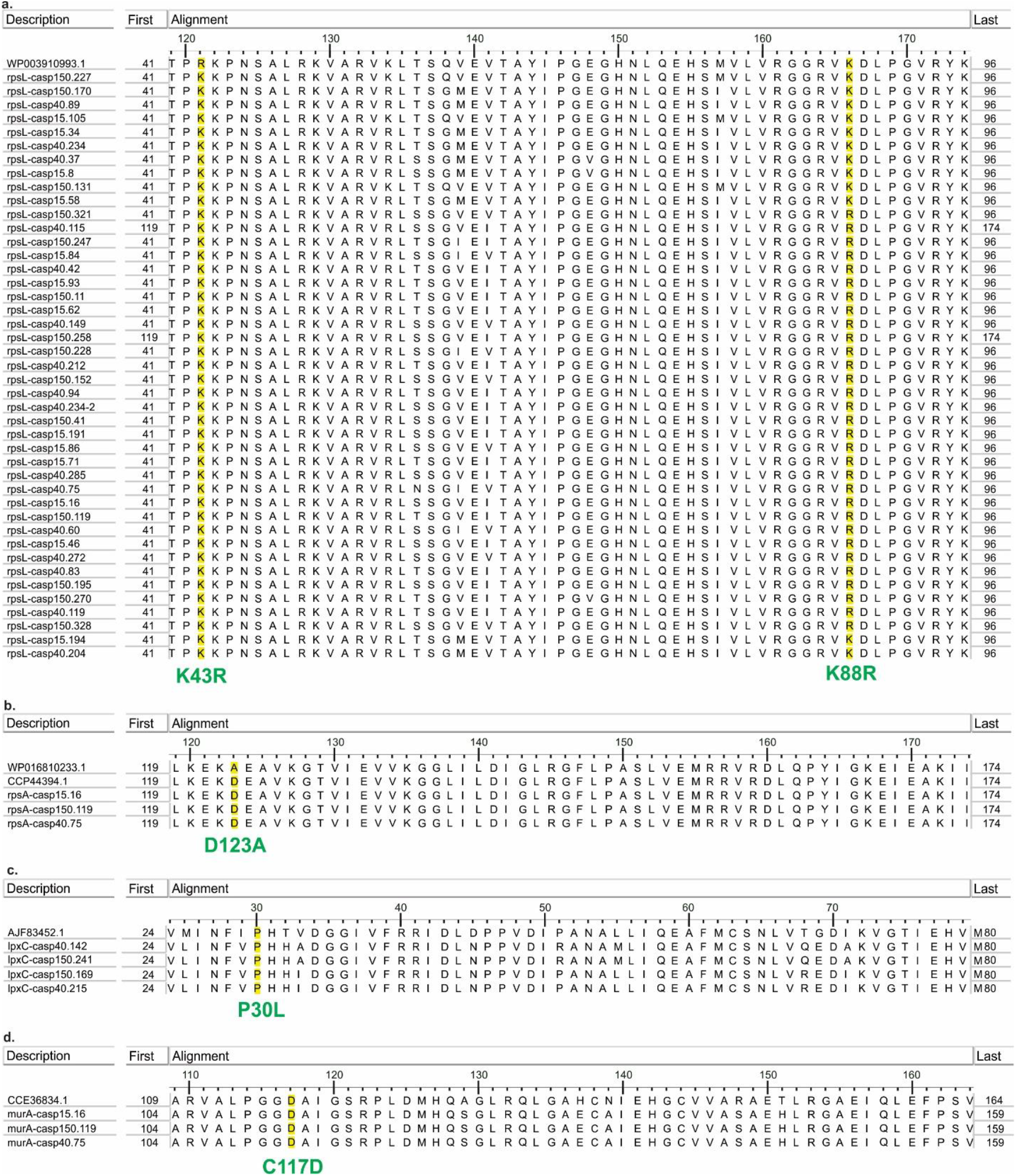
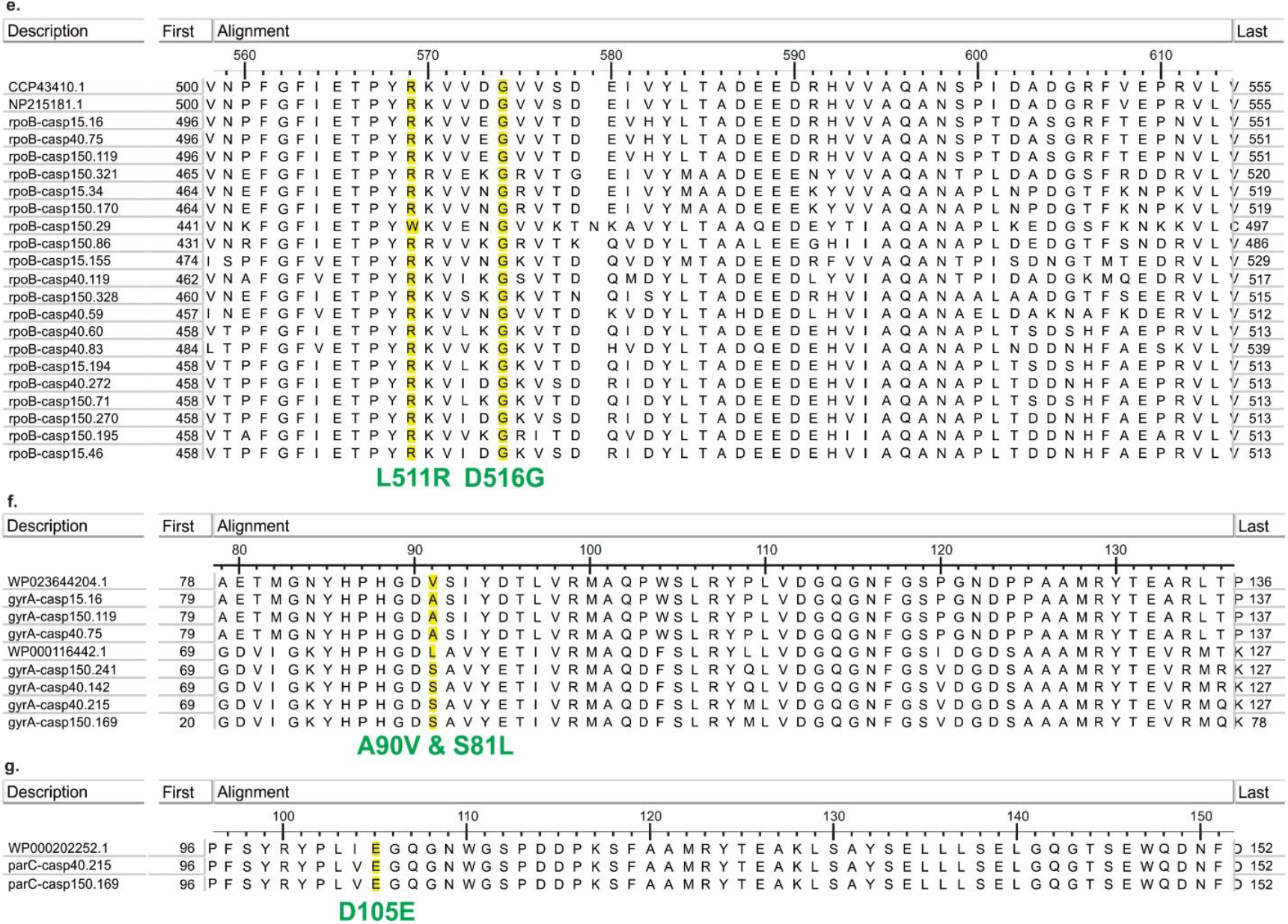
Multiple sequence alignment shows the point mutations that confer the antibiotic resistance. Sequences containing the relevant mutation were confirmed as antibiotic resistance genes. All alignments results are accompanying this manuscript as **Supplementary Data S2**. Reference genes belong to (**a**,**b**,**d**,**e**,**f**) *Mycobacterium tuberculosis* (WP_003910993, WP_016810233, CCP44394, CCE36834, CCP43410, NP_215181, and WP_023644204) and (**c**,**f**,**g**) *Acinetobacter baumannii* (AJF83452, WP_000116442 and WP_000202252). Alignments figures were obtained with the NCBI Genome Workbench v3.7.1.

**Figure S3.**
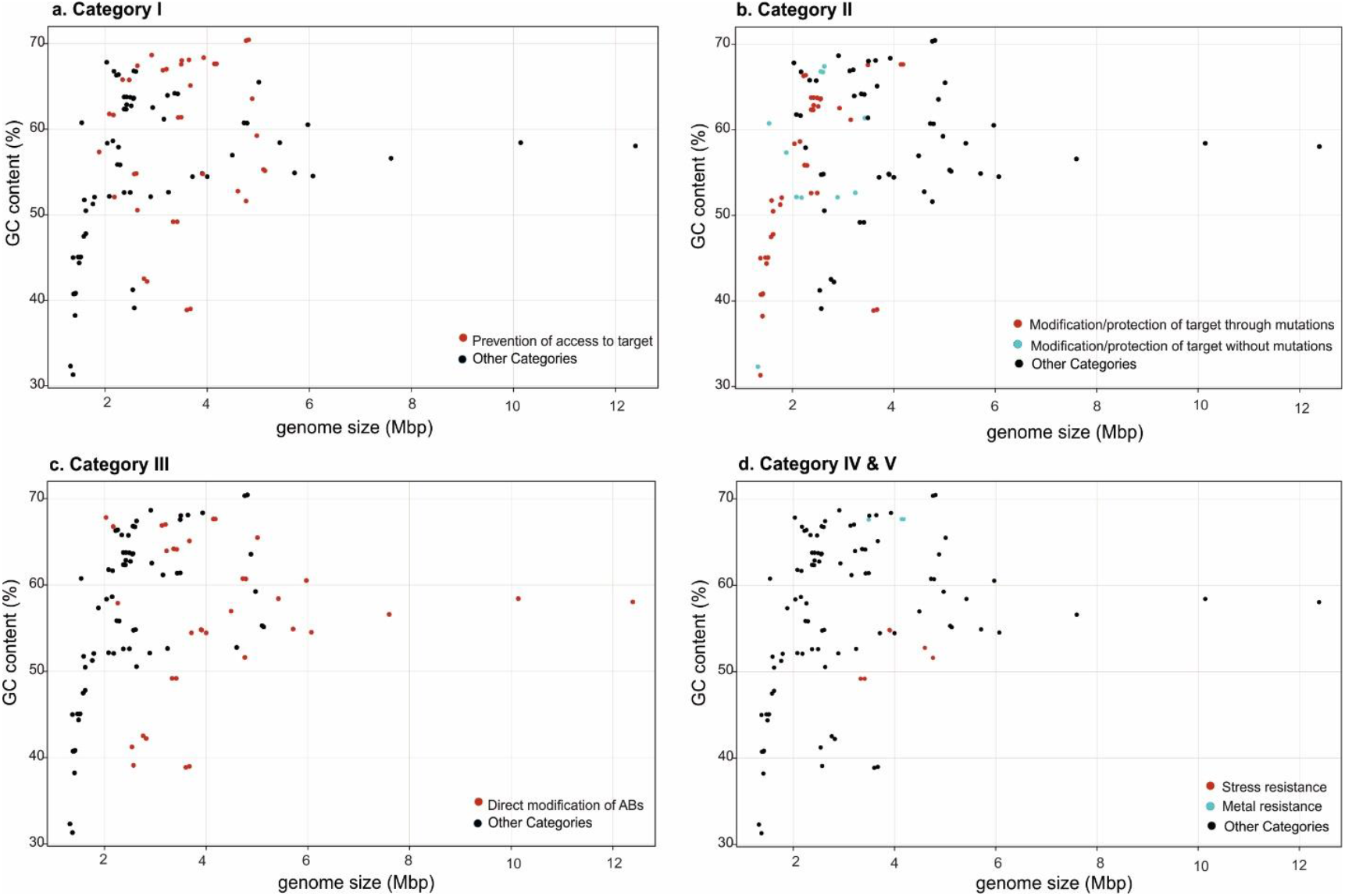
Genomic GC content versus estimated genome size for all ARG containing MAGs. the ARG categories are marked in each plot. Genomic GC content versus estimated genome size are colored based on: (a) Category I, (b)category II (red dots indicate genomes that confer resistant through mutations, which have lower GC content and estimated genome size. Blue dots indicate genomes that confer resistant without mutation.), (c) category III and (d) category IV and V.

**Figure S4.**
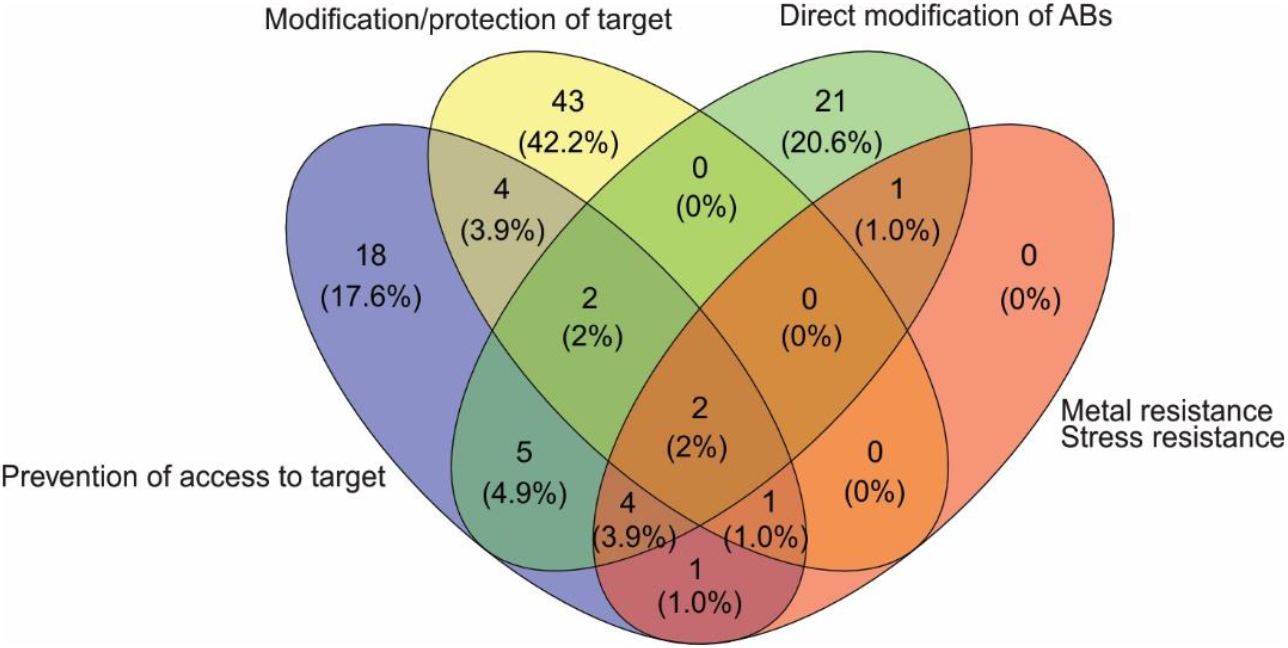
Venn diagram of antibiotic resistance categories. This picture indicates the number of Caspian ARG containing MAGs for each ARG category. 2 MAGs are shared in all the groups and 2 MAGs are shared in three main categories, meaning that these MAGs have various ARGs from diverse categories. Due to the small number of genes in the two categories of Metal resistance and Stress resistance, and also to avoid complicating the Venn diagram, we have considered these two categories as one group.

**Figure S5.**
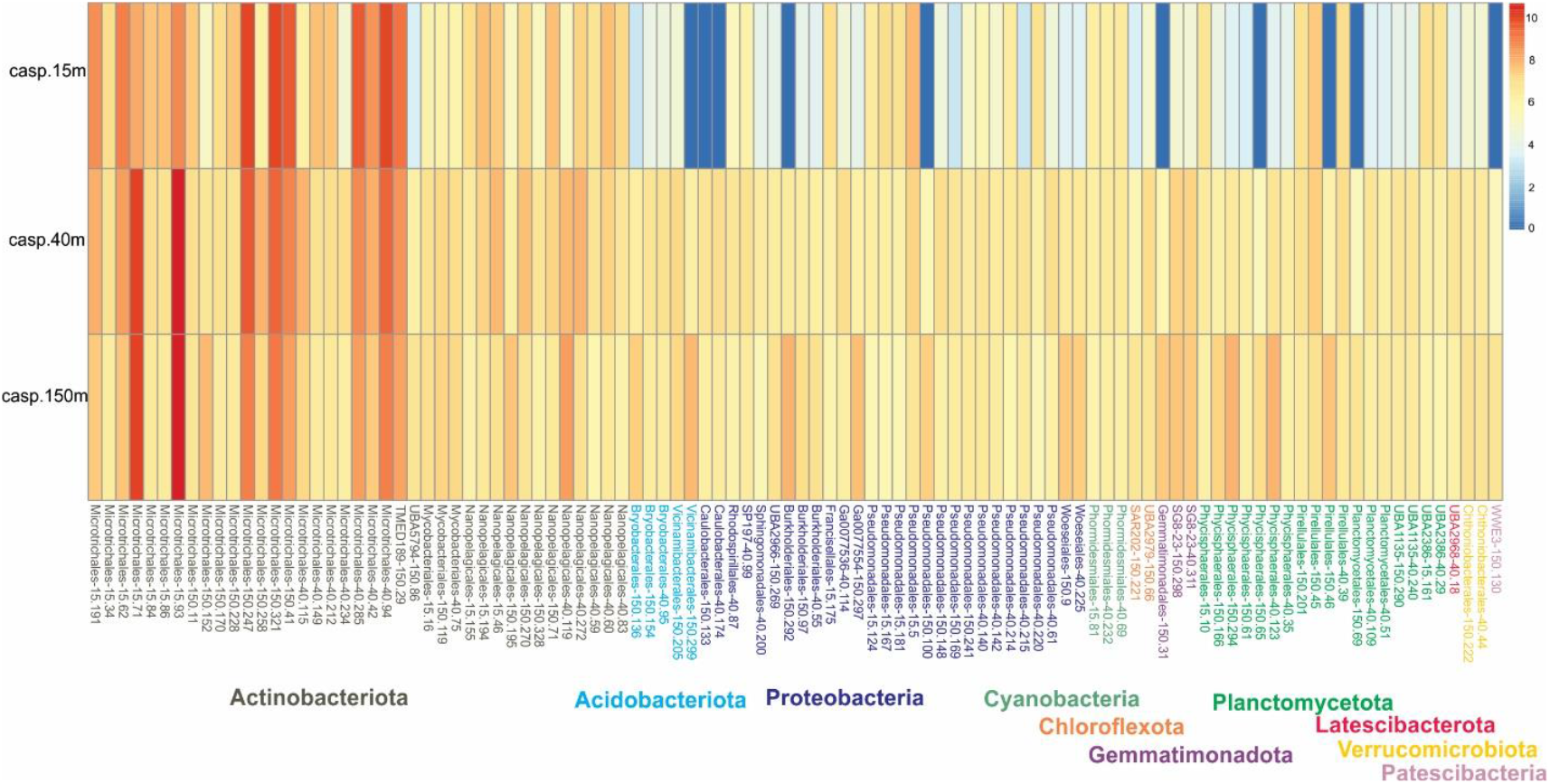
Heat map representation of genome abundance of ARG containing MAGs in different depth of the Caspian Sea. Heat map is drawn in logarithmic scale for better visualizing differences. The members

**Figure S6.**
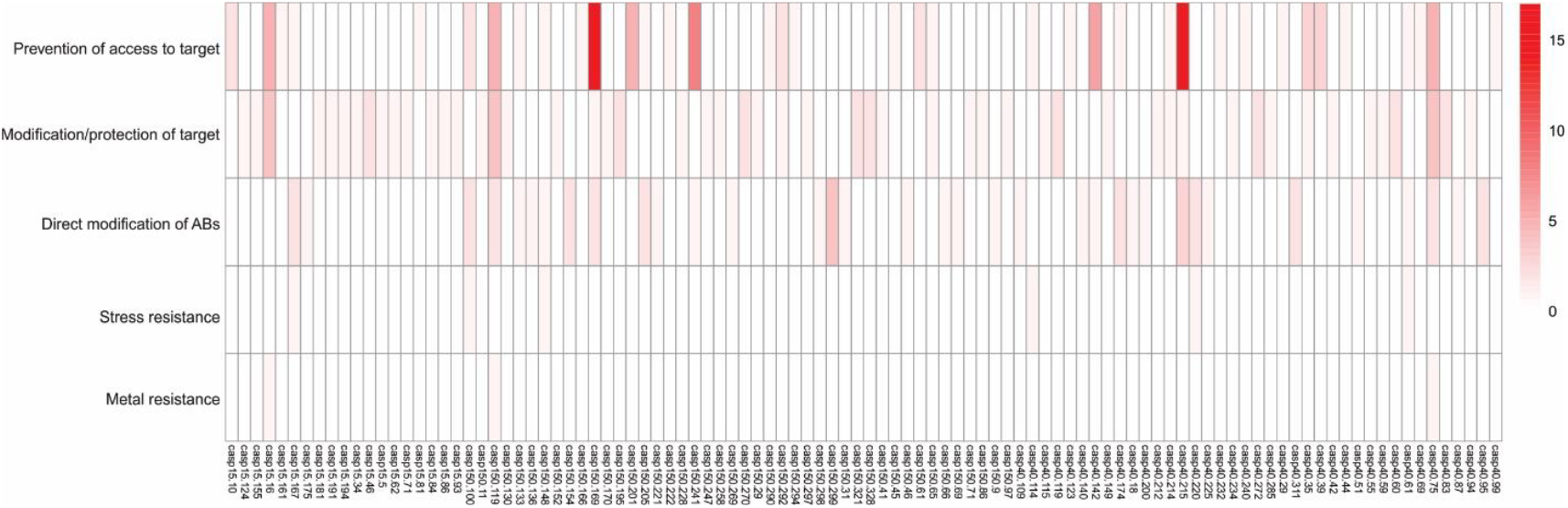
Heat map representation of number of observed ARGs in each Caspian ARG containing MAG with respect to different categories of antibiotic resistance.

**Figure S7.**
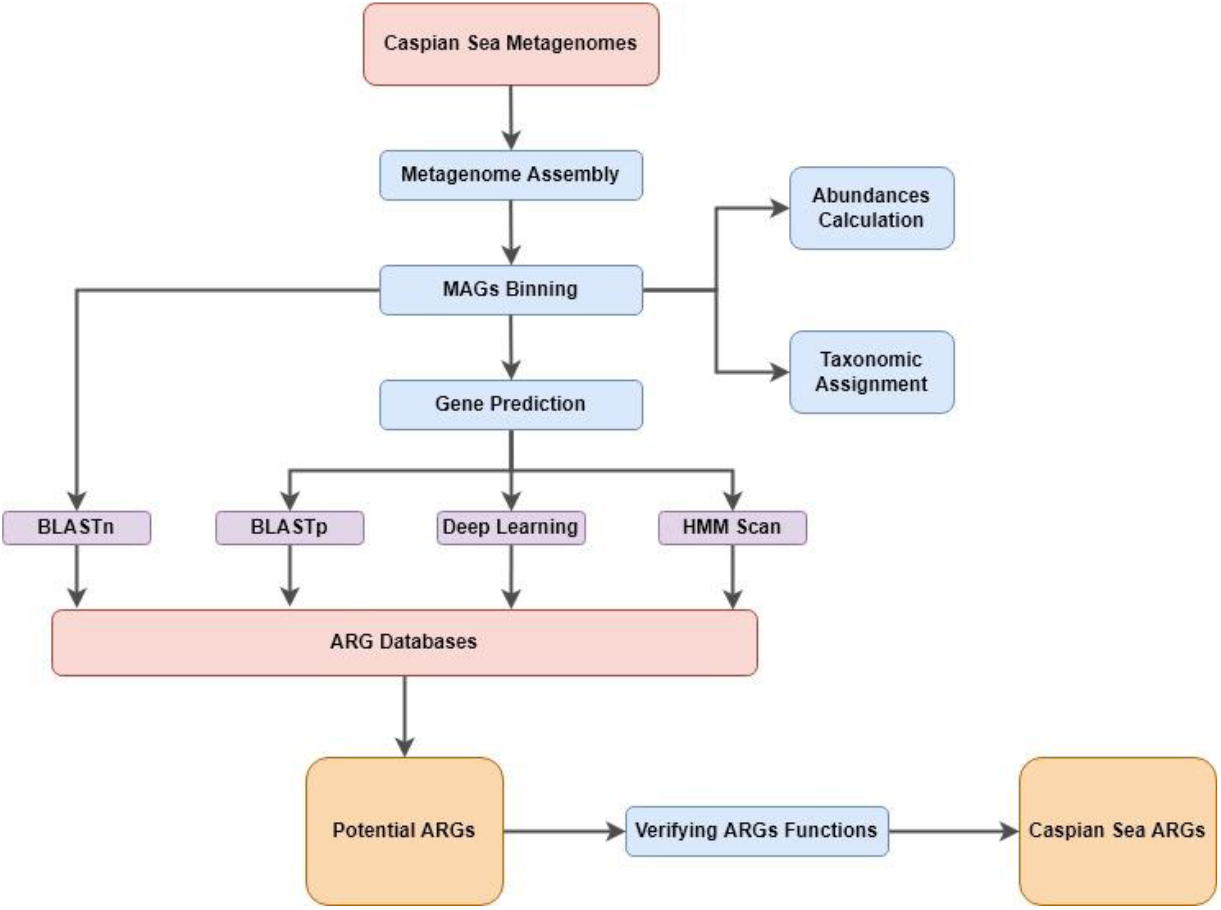
A pipeline workflow diagram describing the steps involved in the Caspian Sea ARGs identification. The red boxes denote inputs, blue boxes represent steps in the study, purple boxes represent the approaches, yellow boxes denote outputs and arrows show the directionality of the workflow.

## References

UN Interagency Coordination Group (IACG) on Antimicrobial Resistance. No Time to Wait: Securing the future from drug-resistant infections. World Heal Organ [Internet]. 2019; Available from: https://www.who.int/antimicrobial-resistance/interagency-coordination-group/IACG_final_report_EN.pdf

Hernando-Amado S, Coque TM, Baquero F, Martínez JL. Defining and combating antibiotic resistance from One Health and Global Health perspectives. Nat Microbiol. 2019;4(9):1432–42.

Sengupta S, Chattopadhyay MK, Grossart H-P. The multifaceted roles of antibiotics and antibiotic resistance in nature. Front Microbiol. 2013;4:47.

Allen HK, Donato J, Wang HH, Cloud-Hansen KA, Davies J, Handelsman J. Call of the wild: antibiotic resistance genes in natural environments. Nat Rev Microbiol. 2010;8(4):251–9.

Hoffman LR, D’Argenio DA, MacCoss MJ, Zhang Z, Jones RA, Miller SI. Aminoglycoside antibiotics induce bacterial biofilm formation. Nature. 2005;436(7054):1171–5.

Skindersoe ME, Alhede M, Phipps R, Yang L, Jensen PO, Rasmussen TB, et al. Effects of antibiotics on quorum sensing in Pseudomonas aeruginosa. Antimicrob Agents Chemother. 2008;52(10):3648–63.

Martinez JL, Sánchez MB, Martínez-Solano L, Hernandez A, Garmendia L, Fajardo A, et al. Functional role of bacterial multidrug efflux pumps in microbial natural ecosystems. FEMS Microbiol Rev. 2009;33(2):430–49.

Aminov RI. The role of antibiotics and antibiotic resistance in nature. Environ Microbiol. 2009;11(12):2970–88.

Larsson DG, Flach C-F. Antibiotic resistance in the environment. Nat Rev Microbiol. 2021;1–13.

Forsberg KJ, Reyes A, Wang B, Selleck EM, Sommer MOA, Dantas G. The shared antibiotic resistome of soil bacteria and human pathogens. Science (80-). 2012;337(6098):1107–11.

Ventola CL. The antibiotic resistance crisis: part 1: causes and threats. Pharm Ther. 2015;40(4):277.

Serwecińska L. Antimicrobials and antibiotic-resistant bacteria: a risk to the environment and to public health. Water. 2020;12(12):3313.

MacFadden DR, McGough SF, Fisman D, Santillana M, Brownstein JS. Antibiotic resistance increases with local temperature. Nat Clim Chang. 2018;8(6):510–4.

Cuadrat RRC, Sorokina M, Andrade BG, Goris T, Davila AMR. Global ocean resistome revealed: Exploring antibiotic resistance gene abundance and distribution in TARA Oceans samples. Gigascience. 2020;9(5):giaa046.

Hatosy SM, Martiny AC. The ocean as a global reservoir of antibiotic resistance genes. Appl Environ Microbiol. 2015;81(21):7593–9.

Moon K, Jeon JH, Kang I, Park KS, Lee K, Cha C-J, et al. Freshwater viral metagenome reveals novel and functional phage-borne antibiotic resistance genes. Microbiome. 2020;8:1–15.

Spänig S, Eick L, Nuy JK, Beisser D, Ip M, Heider D, et al. A multi-omics study on quantifying antimicrobial resistance in European freshwater lakes. Environ Int. 2021;157:106821.

Yang Y, Li Z, Song W, Du L, Ye C, Zhao B, et al. Metagenomic insights into the abundance and composition of resistance genes in aquatic environments: Influence of stratification and geography. Environ Int. 2019;127:371–80.

Zhang H, Wang Y, Liu P, Sun Y, Dong X, Hu X. Unveiling the occurrence, hosts and mobility potential of antibiotic resistance genes in the deep ocean. Sci Total Environ. 2021;151539.

Naddafi R, Koupayeh NH, Ghorbani R. Spatial and temporal variations in stable isotope values (δ13C and δ15N) of the primary and secondary consumers along the southern coastline of the Caspian Sea. Mar Pollut Bull. 2021;164:112001.

WHO report on surveillance of antibiotic consumption: 2016-2018 early implementation. WHO [Internet]. 2018; Available from: https://www.who.int/medicines/areas/rational_use/oms-amr-amc-report-2016-2018/en/

Boolchandani M, D’Souza AW, Dantas G. Sequencing-based methods and resources to study antimicrobial resistance. Nat Rev Genet. 2019;20(6):356–70.

McDermott PF, Tyson GH, Kabera C, Chen Y, Li C, Folster JP, et al. Whole-genome sequencing for detecting antimicrobial resistance in nontyphoidal Salmonella. Antimicrob Agents Chemother. 2016;60(9):5515–20.

Suzuki S, Horinouchi T, Furusawa C. Prediction of antibiotic resistance by gene expression profiles. Nat Commun. 2014;5(1):1–12.

Rodwell TC, Valafar F, Douglas J, Qian L, Garfein RS, Chawla A, et al. Predicting extensively drug-resistant Mycobacterium tuberculosis phenotypes with genetic mutations. J Clin Microbiol. 2014;52(3):781–9.

Mehrshad M, Amoozegar MA, Ghai R, Shahzadeh Fazeli SA, Rodriguez-Valera F. Genome reconstruction from metagenomic data sets reveals novel microbes in the brackish waters of the Caspian Sea. Appl Environ Microbiol. 2016;82(5):1599–612.

Baharoglu Z, Garriss G, Mazel D. Multiple pathways of genome plasticity leading to development of antibiotic resistance. Antibiotics. 2013;2(2):288–315.

Di Cesare A, Sabatino R, Yang Y, Brambilla D, Li P, Fontaneto D, et al. Contribution of plasmidome, metal resistome and integrases to the persistence of the antibiotic resistome in aquatic environments. Environ Pollut. 2021;118774.

Khare G, Nangpal P, Tyagi AK. Differential roles of iron storage proteins in maintaining the iron homeostasis in Mycobacterium tuberculosis. PLoS One. 2017;12(1):e0169545.

Bereswill S, Waidner U, Odenbreit S, Lichte F, Fassbinder F, Kist M. Structural, functional and mutational analysis of the pfr gene encoding a ferritin from Helicobacter pylori. Microbiology. 1998;144(9):2505–16.

Dantas G, Sommer MOA, Oluwasegun RD, Church GM. Bacteria subsisting on antibiotics. Science (80-). 2008;320(5872):100–3.

Crofts TS, Wang B, Spivak A, Gianoulis TA, Forsberg KJ, Gibson MK, et al. Shared strategies for β-lactam catabolism in the soil microbiome. Nat Chem Biol. 2018;14(6):556–64.

Morgado SM, Vicente ACP. Comprehensive in silico survey of the Mycolicibacterium mobilome reveals an as yet underexplored diversity. Microb genomics. 2021;7(3).

Komatsu T, Ohya K, Sawai K, Odoi JO, Otsu K, Ota A, et al. Draft genome sequences of Mycolicibacterium peregrinum isolated from a pig with lymphadenitis and from soil on the same Japanese pig farm. BMC Res Notes. 2019;12(1):1–4.

Baumann P. Isolation of Acinetobacter from soil and water. J Bacteriol. 1968;96(1):39–42.

Lee C-R, Lee JH, Park M, Park KS, Bae IK, Kim YB, et al. Biology of Acinetobacter baumannii: pathogenesis, antibiotic resistance mechanisms, and prospective treatment options. Front Cell Infect Microbiol. 2017;7:55.

Neuenschwander SM, Ghai R, Pernthaler J, Salcher MM. Microdiversification in genome-streamlined ubiquitous freshwater Actinobacteria. ISME J. 2018;12(1):185–98.

Giovannoni SJ, Thrash JC, Temperton B. Implications of streamlining theory for microbial ecology. ISME J. 2014;8(8):1553–65.

Farhangi MB, Ghorbanzadeh N, Amini M, Ghovvati S. Investigation of antibiotic resistant coliform bacteria in Zarjoub River. Iran J Soil Water Res. 2021;52(8):2061–76.

Saberinia F, Farhangi MB, Yaghmaeian Mahabadi N, Ghorbanzadeh N. Investigation of Gowharrood River Contamination to Antibiotic Resistant Bacteria. J Water Wastewater; Ab va Fazilab (in persian). 2021;31(7):145–61.

Bush K, Jacoby GA. Updated functional classification of β-lactamases. Antimicrob Agents Chemother. 2010;54(3):969–76.

Boyd SE, Livermore DM, Hooper DC, Hope WW. Metallo-β-lactamases: structure, function, epidemiology, treatment options, and the development pipeline. Antimicrob Agents Chemother. 2020;64(10):e00397–20.

Gatica J, Jurkevitch E, Cytryn E. Comparative metagenomics and network analyses provide novel insights into the scope and distribution of β-lactamase homologs in the environment. Front Microbiol. 2019;10:146.

Hofer U. Feasting on β-lactams. Nat Rev Microbiol. 2018;16(7):394–5.

Nurk S, Meleshko D, Korobeynikov A, Pevzner PA. metaSPAdes: a new versatile metagenomic assembler. Genome Res. 2017;27(5):824–34.

Kang DD, Li F, Kirton E, Thomas A, Egan R, An H, et al. MetaBAT 2: an adaptive binning algorithm for robust and efficient genome reconstruction from metagenome assemblies. PeerJ. 2019;7:e7359.

Parks DH, Imelfort M, Skennerton CT, Hugenholtz P, Tyson GW. CheckM: assessing the quality of microbial genomes recovered from isolates, single cells, and metagenomes. Genome Res. 2015;25(7):1043–55.

Chaumeil P-A, Mussig AJ, Hugenholtz P, Parks DH. GTDB-Tk: a toolkit to classify genomes with the Genome Taxonomy Database. Oxford University Press; 2020.

Hyatt D, Chen G-L, LoCascio PF, Land ML, Larimer FW, Hauser LJ. Prodigal: prokaryotic gene recognition and translation initiation site identification. BMC Bioinformatics. 2010;11(1):1–11.

Alcock BP, Raphenya AR, Lau TTY, Tsang KK, Bouchard M, Edalatmand A, et al. CARD 2020: antibiotic resistome surveillance with the comprehensive antibiotic resistance database. Nucleic Acids Res [Internet]. 2020 Jan 8;48(D1):D517–25. Available from: https://doi.org/10.1093/nar/gkz935

Feldgarden M, Brover V, Haft DH, Prasad AB, Slotta DJ, Tolstoy I, et al. Validating the AMRFinder tool and resistance gene database by using antimicrobial resistance genotype-phenotype correlations in a collection of isolates. Antimicrob Agents Chemother. 2019;63(11):e00483–19.

Arango-Argoty G, Garner E, Pruden A, Heath LS, Vikesland P, Zhang L. DeepARG: a deep learning approach for predicting antibiotic resistance genes from metagenomic data. Microbiome [Internet]. 2018;6(1):23. Available from: https://doi.org/10.1186/s40168-018-0401-z

Bortolaia V, Kaas RS, Ruppe E, Roberts MC, Schwarz S, Cattoir V, et al. ResFinder 4.0 for predictions of phenotypes from genotypes. J Antimicrob Chemother [Internet]. 2020 Dec 1;75(12):3491–500. Available from: https://doi.org/10.1093/jac/dkaa345

Arango-Argoty GA, Guron GKP, Garner E, Riquelme M V, Heath LS, Pruden A, et al. ARGminer: a web platform for the crowdsourcing-based curation of antibiotic resistance genes. Bioinformatics. 2020;36(9):2966–73.

Pal C, Bengtsson-Palme J, Rensing C, Kristiansson E, Larsson DGJ. BacMet: antibacterial biocide and metal resistance genes database. Nucleic Acids Res. 2014;42(D1):D737–43.

Panunzi LG. sraX: A Novel Comprehensive Resistome Analysis Tool [Internet]. Vol. 11, Frontiers in Microbiology. 2020. p. 52. Available from: https://www.frontiersin.org/article/10.3389/fmicb.2020.00052

Kumar GS, Roshan PB, M. Ds, Rafael L-R, Marie K, Luce L, et al. ARG-ANNOT, a New Bioinformatic Tool To Discover Antibiotic Resistance Genes in Bacterial Genomes. Antimicrob Agents Chemother [Internet]. 2014 Jan 1;58(1):212–20. Available from: https://doi.org/10.1128/AAC.01310-13

Seemann T. Abricate [Internet]. 2019. Available from: https://github.com/tseemann/abricate

Lu S, Wang J, Chitsaz F, Derbyshire MK, Geer RC, Gonzales NR, et al. CDD/SPARCLE: the conserved domain database in 2020. Nucleic Acids Res. 2020;48(D1):D265–8.

Potter SC, Luciani A, Eddy SR, Park Y, Lopez R, Finn RD. HMMER web server: 2018 update. Nucleic Acids Res. 2018;46(W1):W200–4.

Mistry J, Chuguransky S, Williams L, Qureshi M, Salazar GA, Sonnhammer ELL, et al. Pfam: The protein families database in 2021. Nucleic Acids Res [Internet]. 2021 Jan 8;49(D1):D412–9. Available from: https://doi.org/10.1093/nar/gkaa913

Kanehisa M, Sato Y, Morishima K. BlastKOALA and GhostKOALA: KEGG tools for functional characterization of genome and metagenome sequences. J Mol Biol. 2016;428(4):726–31.

Huerta-Cepas J, Szklarczyk D, Heller D, Hernández-Plaza A, Forslund SK, Cook H, et al. eggNOG 5.0: a hierarchical, functionally and phylogenetically annotated orthology resource based on 5090 organisms and 2502 viruses. Nucleic Acids Res. 2019;47(D1):D309–14.

Thompson JD, Higgins DG, Gibson TJ. CLUSTAL W: improving the sensitivity of progressive multiple sequence alignment through sequence weighting, position-specific gap penalties and weight matrix choice. Nucleic Acids Res. 1994;22(22):4673–80.

Kumar S, Stecher G, Li M, Knyaz C, Tamura K. MEGA X: molecular evolutionary genetics analysis across computing platforms. Mol Biol Evol. 2018;35(6):1547.

Letunic I, Bork P. Interactive Tree Of Life (iTOL) v5: an online tool for phylogenetic tree display and annotation. Nucleic Acids Res. 2021;49(W1):W293–6.

Guo J, Bolduc B, Zayed AA, Varsani A, Dominguez-Huerta G, Delmont TO, et al. VirSorter2: a multi-classifier, expert-guided approach to detect diverse DNA and RNA viruses. Microbiome. 2021;9(1):1–13.

